# The cellular architecture of microvessels, pericytes and neuronal cell types in organizing regional brain energy homeostasis in mice

**DOI:** 10.1101/2021.05.19.444854

**Authors:** Yuan-ting Wu, Hannah C. Bennett, Uree Chon, Daniel J. Vanselow, Qingguang Zhang, Rodrigo Muñoz-Castañeda, Keith C. Cheng, Pavel Osten, Patrick J. Drew, Yongsoo Kim

## Abstract

Cerebrovasculature and its mural cells must meet dynamic energy demands of different neuronal cell types across the brain, but their spatial relationship is largely unknown. Here, we apply brain-wide mapping methods to create a comprehensive cellular-resolution resource comprising the distribution of and quantitative relationship between cerebrovasculature, pericytes, and glutamatergic and GABAergic neurons, including neuronal nitric oxide synthase-positive (nNOS+) neurons and their subtypes, as well as simulation-ready vascular tracing data in mice. We discover strikingly high densities of vasculature and pericytes with high blood perfusion in primary motor-sensory cortices compared to association cortices that show significant positive and negative correlation with parvalbumin+ and nNOS+ neurons, respectively. Thalamo-striatal areas linked to primary motor-sensory cortices also contain high densities of vasculature and pericytes compared to association areas. Collectively, our results unveil a finely tuned spatial relationship between cerebrovascular network and neuronal cell composition in meeting regional energy needs of the brain.

## Introduction

The brain is the most energy-demanding organ per gram and is powered by an intricate web of vascular and mural cells that dynamically supply blood and clear metabolic waste ^1–4^. Pericytes, a key mural cell type, wrap around microvessels and are proposed to regulate blood flow and vascular permeability ^2,5–8^. Moreover, neuronal activity is known to regulate vascular diameter directly or indirectly (via astrocytes), which is referred to as a neurovascular coupling ^9,10^. Neurons, unlike muscle cells, strictly rely on aerobic metabolism, thus neuronal functions critically depend on efficient vascular support ^3,11^. Not surprisingly, impairment of the cerebrovasculature, pericytes, and neurovascular coupling has been widely implicated in many neurological disorders such as stroke and neurodegenerative disorders ^4^. Yet, despite its significance, we have limited knowledge on the cellular architecture of vasculature and pericytes, especially with respect to their quantitative relationship with neuronal cell types across the whole brain. This relationship is likely of critical importance for the heterogeneous coupling of neural activity to blood flow described across different brain regions ^12–15^.

The generation of action potentials and synaptic transmission are energetically demanding ^16^ and accordingly energy consumption is proposed to be linearly correlated with the number of neurons across different animal species including humans ^17^. However, neurons comprise highly distinct subtypes with different morphological, electrophysiological, and molecular characteristics ^18,19^. For example, the major classes of GABAergic neurons in the cortex include parvalbumin (PV), somatostatin (SST), and vasoactive intestinal peptide (VIP) expressing neurons, each of which make distinct synaptic connections with pyramidal neurons and each other ^18^. Moreover, these neuronal cell types are expressed at different densities across cortical areas: PV interneurons have high density in sensory cortices and low density in association cortices, while SST neurons show the opposite density pattern in mice ^20^. Different neuronal subtypes also have differential energy demands and regulation. For instance, the fast spiking PV neurons are among the highest energy demanding neuronal types ^21^. On the other hand, another neuronal type –neuronal nitric oxide synthase (nNOS) expressing neurons – can actively regulate blood supply by causing vasodilation ^22–24^. Taken together, these data suggest that determining specific spatial relationships between neuronal cell types and the vascular network is critically important for understanding the demand for and the mechanism of distinct blood flow regulation across different brain regions.

It has been technically challenging to comprehensively examine mammalian cerebrovasculature because of its fine and complex branching across the entire brain, with the diameter of small capillaries being only ~5μm. However, recent progress in meso-scale mapping methods began elucidating the structural organization of the cerebrovasculature and its regional heterogeneity across the mouse brain at unprecedented detail ^25–28^. While these studies provide a detailed description of the general organization of mouse brain cerebrovasculature, the question of how cerebrovasculature and its mural cells are structurally organized to support different neuronal cell types across the whole brain remains largely unanswered. To address the relationship between cerebrovasculature and mural and neuronal cell types, we have devised a serial two-photon tomography (STPT)-based imaging approach with newly developed analytical tools to derive the first cellular architecture atlas containing cerebrovasculature, capillary pericytes, and several major neuronal cell types, including PV interneurons and vasomotor nNOS neurons in the adult mouse brain.

Leveraging this data resource, we uncovered several organizational principles of the brain. First, primary motor-sensory cortical areas and related thalamic and dorsal striatal areas have higher densities of vasculature and pericytes compared to association cortices and related thalamo-cortico-striatal pathways. Second, our computational simulation showed that cerebrovasculature in the primary motor-sensory cortices is structured to provide higher blood perfusion compared to association cortices. Third, we found a striking positive correlation between vascular and pericyte densities with energy demanding PV interneurons, but a negative correlation with vasomotor nNOS neurons in the isocortex. Lastly, we present web-based and freely downloadable datasets for the vasculature, pericytes, and 10 different neuronal cell types in the whole mouse brain as a comprehensive resource to facilitate future investigations on understanding brain energy homeostasis in largely under-studied subcortical areas.

## Results

### Comprehensive vascular mapping in the intact mouse brain

Our first goal was to examine the spatial arrangement of the cerebrovasculature in a whole intact mouse brain to understand the anatomical basis of the vascular network. We filled microvessels from 2-month-old C57BL/6 mice with a cardiac perfusion of fluorescein isothiocyanate (FITC)-conjugated albumin gel (Figure 1a) ^29,30^ and used STPT imaging in combination with two-photon optical scans and serial sectioning to image the whole mouse brain at 1×1×5 μm (*x,y,z;* media-lateral, dorsal-ventral, rostral-caudal) (Figure 1b). Since the vascular tracing requires precise stitching across X-Y tiles throughout z stacks, we developed a new stitching algorithm to correct optical aberrations, bleaching in overlapped tile areas, and z stack alignments (Figure 1c-d; Figure S1). To visualize and quantitatively analyze cerebrovascular arrangement, we developed a computational pipeline that binarizes and traces the original image of the FITC-filled vasculature, as well as quantifies the diameter of each vessel and its branching points across the whole brain (Figure 1e-g; See Methods for more details). Individual brains were then registered to the Allen Common Coordinate Framework (CCF) to quantify signals (vascular density, branching density, and vascular radii) across different anatomical areas ^31^ (Figure 1h-l; Table S1; Movie S1). Although we observed near-complete vasculature labeling with our FITC-based filling approach, we implemented an additional quality control step to reject data from areas with potentially incomplete labeling or imaging artifacts (Figure S1). Moreover, we confirmed that our approach closely reflects vasculature *in vivo* by directly comparing STPT results with *in vivo* two-photon measurements of the same vasculature acquired in the same mice (Figure S2).

**Figure 1.**
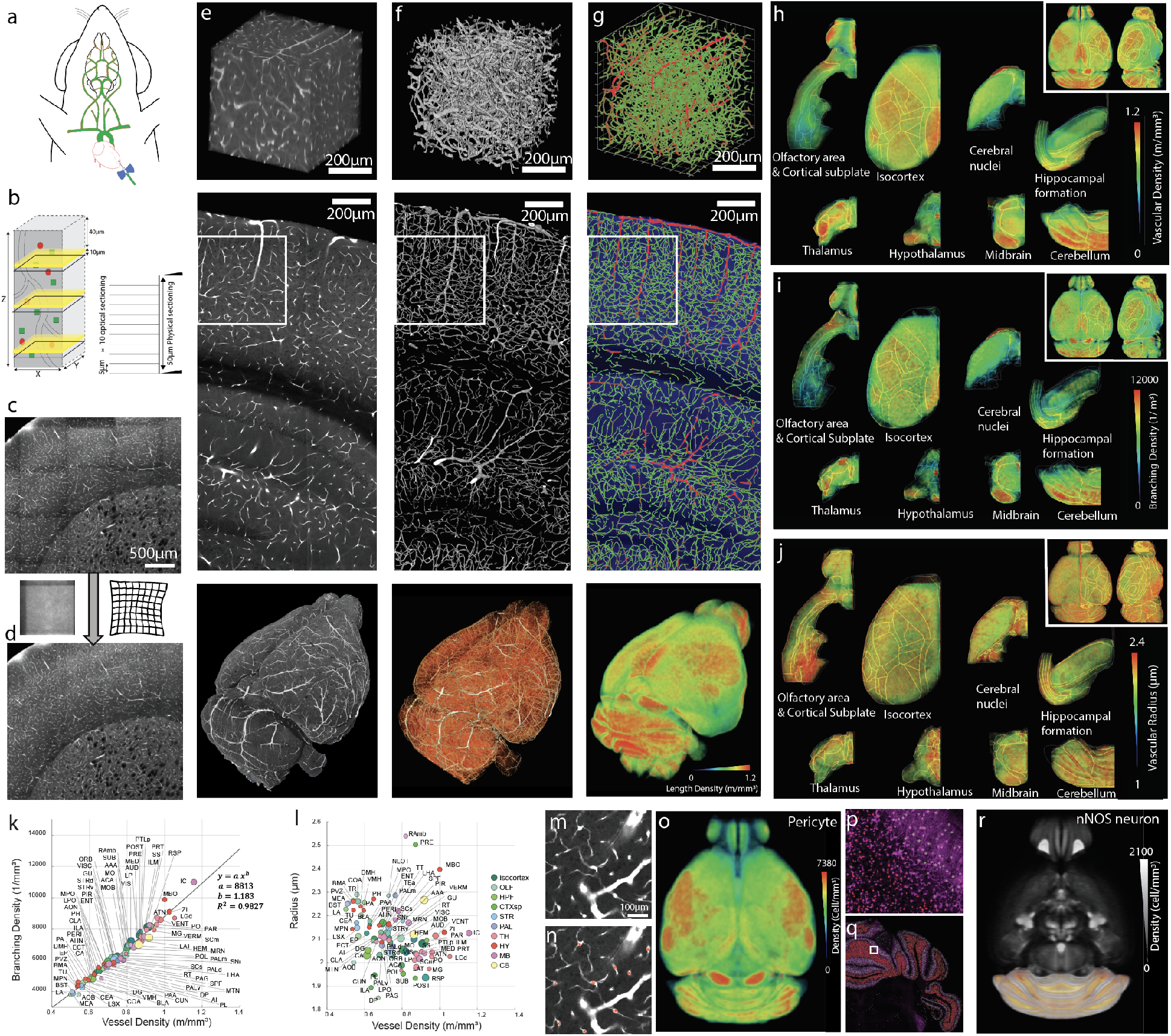
High resolution 3D mapping of the cerebrovasculature, pericytes, and neuronal cell types. **a**. Fluorescent dye (FITC)-conjugated albumin gel perfusing the mouse brain through the heart to label cerebrovasculature. **b**. Combination of physical sectioning (vibratome cutting) and optical sectioning to achieve lossless imaging of a sample. Left: black-plane indicates the physical cuts, yellow-zones indicates an optical sectioning imaging plane. Right: 10 optical imaging sections per one physical sectioning. **c-d**. Stitching with optical aberration and tile line correction (**d**) from uncorrected images (**c**). **e-g**. Example outputs from each stage of the analysis pipeline. Top row: 100 μm thick 3D volume from the white box areas from the middle row, Middle row: An example coronal section, Bottom row: The whole brain results. **e.** The raw image volume of FITC labeled vasculature. **f**. The binarized vasculature. **g**. The traced vasculature. Large (radius > 5 μm) and small vessels are colored as red and green, respectively in the top and middle image. The bottom image shows the vasculature density. **h-j**. The averaged vasculature length density (**h**), branching density (**i**), and radii (**j**) from four brains are registered to the Allen CCF and displayed with heatmap in 8 major areas across the whole mouse brain. **k-l**. The correlation between vessel density and branching density (**k**) and the correlation between vessel density and the averaged radius (**l**). Size of each ROI is displayed according to relative volume of the area. Note strong correlation between vessel density with branching density, but not with vascular radii. Please see Table S1 for abbreviations. **m-o.** Pericyte density mapping. Example of tdTomato labeling from PDGFRβ-Cre:Ai14 mice (**m**) and pericyte detection algorithm (red stars, **n**). Brain-wide pericyte density (**o**). **p-r**. pan nNOS neuronal mapping using nNOS-CreER:Ai14 mouse. AI based detection of nNOS cells with two distinct shapes (green and red crosses) in the cerebellum (**p** from the white box in **q**). Brain-wide nNOS density (**r**).

Having established a precise methodology for complete vasculature mapping, our second goal was to map the density distribution of pericytes and neuronal nitric oxide synthase (nNOS) expressing neurons as vasomodulatory cell types, and major cortical cell types (pan glutamatergic, pan GABAergic, PV, SST, and VIP neurons) with different energy demands in order to understand quantitative spatial relationship between these cell types and cerebrovasculature (Table 1). We employed a genetic labeling method using cell type specific Cre drivers crossed with conditional reporter mice to label mural cell types including pericytes using platelet-derived growth factor receptor beta (PDGFRβ)-Cre:Ai14 mice ^1,32^ and neuronal cell types including nNOS neurons ^33^ (Table 1). Moreover, we utilized genetic intersection methods using Cre and Flp dependent reporter mice (Ai65) crossed with nNOS-CreER and one of the subtype markers (Neuropeptide Y;NPY-, SST-, PV-, or VIP-Flp) to label nNOS subtypes linked with distinct functions including regulating vascular diameter ^34,35^ (Table 1). To conduct whole brain quantification of these cell populations after STPT imaging, we developed new deep learning algorithms to specifically count capillary pericytes from fluorescently-labeled cells in PDGFRβ-Cre:Ai14 mice and to detect neurons including densely packed nNOS neurons in the cerebellum (Figure 1m-n, p-q; Figure S3; Table S2; see the methods section for more details). Detected signals were then registered onto the adult CCF to quantify the 3D density of the target cell type distribution across the whole brain (Kim et al., 2017; Wang et al., 2020) (Figure 1o,r; Table S3-S5; Movie S2-S4).

**Table 1:** Cell type specific transgenic mouse lines to label mural and neuronal cell types.

| Cell Type | Transgenic mouse lines |
| --- | --- |
| Mural cell types including Pericyte | PDGFRb-Cre:Ai14 |
| pan nNOS neurons | nNOS-CreER:Ai14 |
| nNOS subtype co-expressing NPY | nNOS-CreER:NPY-Flp:Ai65 |
| nNOS subtype co-expressing SST | nNOS-CreER:SST-Flp:Ai65 |
| nNOS subtype co-expressing PV | nNOS-CreER:PV-Flp:Ai65 |
| nNOS subtype co-expressing VIP | nNOS-CreER:VIP-Flp:Ai65 |
| pan glutamatergic neurons | vGlut1-Cre:Ai75 |
| pan GABAergic neurons | Gad2-Cre:Ai75 |
| GABAergic subtype expressing PV | PV-Cre:H2B-GFP |
| GABAergic subtype expressing SST | SST-Cre:H2B-GFP |
| GABAergic subtype expressing VIP | VIP-Cre:H2B-GFP |

### Vasculature across the isocortex is not uniform

We focused our analysis first on the isocortex. The mouse isocortex, particularly the primary somatosensory cortex, has been a major area of focus for *in vivo* studies examining the cellular mechanisms of neurovascular coupling ^7,22,23,36^. Yet, it is not known whether the vasculature is anatomically organized in a similar way across the entire cortex or whether different cortical areas may have unique vascular organization signatures and consequently different neurovascular coupling ^13,15^.

To examine the spatial distribution intuitively while maintaining high resolution information, we devised an isocortical flatmap based on Laplace’s equation (Figure 2a-d; see Methods for more details). We grouped isocortical areas into 5 subregions based on their anatomical connectivity and cell type composition; motor somatosensory, audio visual, medial association, medial prefrontal, and lateral association ^20,37^ (Figure 2d). When averaged vessel length density is plotted in the cortical flatmap, it is clear that primary motor and sensory (auditory, somatosensory, visual) cortices contain higher vascular densities than association areas (medial prefrontal, medial and lateral association) (Figure 2d-h). For example, densely vascularized areas are tightly aligned with anatomical borders in the somatosensory (SS) and primary auditory (AUDp) cortices (Figure 2e, grey and white arrowheads). One notable exception is the ventral retrosplenial (RSPv) cortex, a part of the medial association group, that receive spatial navigation information from the dorsal subiculum ^37^ and shows remarkably high vascular density compared to other cortical areas (Figure 2e, black arrowhead). Cortical layer-specific maximum projection of the length density shows that sharp boundaries between cortical areas are strongly driven by the layer 2/3/4 (specifically the layer 4 for primary sensory regions) vascular distribution (Figure 2h).

**Figure 2.**
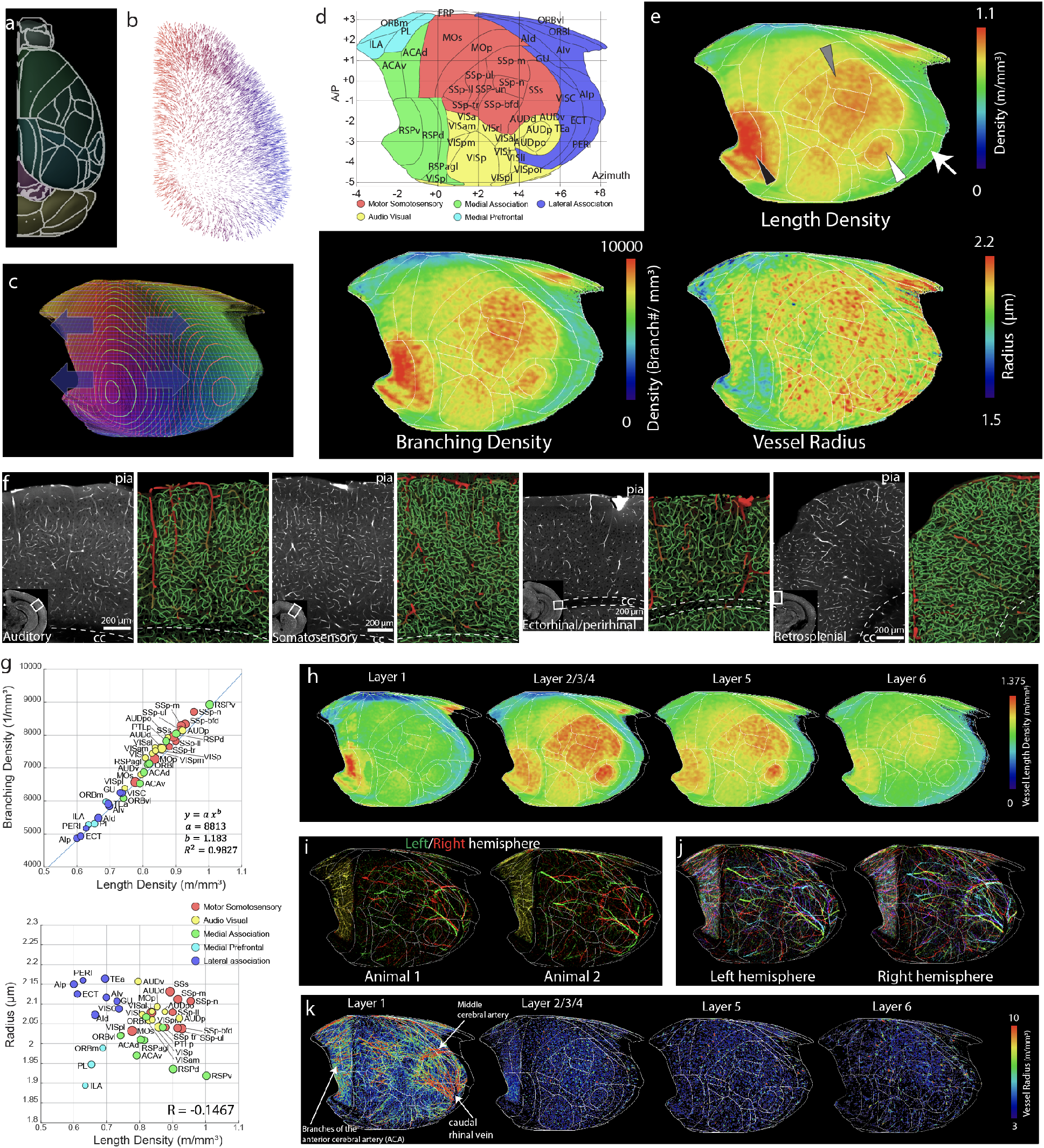
Heterogeneous vascular arrangements in the isocortex. **a-c.** Creating an isocortical flatmap. **a.** Anatomical borderlines of the Allen CCF. **b**. Gradient vectors from solving the Laplace equation by setting cortical layer 1 and layer 6 as end points. **c**. The flattened projected profile. Each line is 0.2 mm apart. The four arrowheads indicate the flattening direction while maintaining the original A-P coordinates. **d**. The cortical flatmap with Allen CCF border line. Y axis: Bregma anterior-posterior (A-P) coordinates, × axis: Azimuth coordinate represents the physical distance by tracing the cortical surface on the coronal cut. **e**. The averaged vasculature length and branching density as well as vessel radius plotted onto the cortical flat map. Note the high density of vasculature in the somatosensory (gray arrowhead), auditory (white arrowhead), and retrosplenial (black arrowhead) cortex, while there is a low density in lateral association cortex (white arrow). **f**. Examples of cortical areas with different vasculature structures. Large (radius > 5 μm) and small vessels are colored as red and green, respectively. **g**. Correlation between average vessel density and branching density (top), or average radius (bottom) in the isocortex. See Table S1 for abbreviation. **h**. Cortical layer specific max projection of vasculature length density. **i**. large surface vessels from left (green) and right (red) hemisphere from two different animals. **j**. large surface vessels from 4 different samples with different colors (red, green, blue, and cyan) in each hemisphere. **k**. large surface and penetrating vessels (8 hemispheres from 4 animals) in cortical layer specific flatmap based on their radius.

Next we analyzed vascular branching density and radius. Branching density closely followed the pattern of the vessel length density (Figure 2e-g). In contrast, average vessel radius did not show significant correlation to the vessel length density (Figure 2e-g). However, plotting vessel radius against vessel length densities unveiled distinct patterns between the five cortical groups. For instance, motor sensory areas showed overall high vascular density and radii while the medial prefrontal group were low in both measurements (Figure 2g). Moreover, the lateral association group showed high vascular radius with relatively lower vascular density and the medial association regions showed the opposite pattern (Figure 2g). Noticeably, the RSPv showed overall low average vascular radius despite a high vascular length density (Figure 2e-g).

Considering that our vascular measurements are largely driven by microvessels due to their abundance, we next examined whether relatively large vasculature (radius ≥ 3 μm) also showed regional variabilities. First, we examined whether surface vasculature is stereotypically organized between samples and even between hemispheres of the same brain. We observed that the positions of large surface vasculatures differ considerably between brains and even between hemispheres from the same brain (Figure 2i-j). When we plotted distribution of the large vasculature from 8 hemispheres from 4 animals in our layer specific flatmap, the layer 1 map including the surface vessel showed that the large vessels including the middle cerebral artery (MCA) closely surround the somatosensory area (Figure 2k, layer 1). Moreover, high density of large penetrating vasculatures was clearly observed in the primary motor sensory areas and the RSPv in the layer 2-4 with a gradual decrease in deeper layers (Figure 2k).

Taken together, these data provide strong evidence that cortical vascularization is not uniform but is distinctly organized in functionally different cortical areas. Most notably, areas processing primary motor-sensory regions, including the RSPv for navigational information, have hypervascularization while integrative association areas show relatively low vascular densities. This suggests that the metabolic demand of rapid signal processing in primary motor-sensory areas is accompanied by a dense vascular network compared to cortical areas involved in more downstream information processing.

### Quantification of vascular permeability reveals spatial heterogeneity of blood perfusion

Microvessels provide a large surface area to supply energy sources (e.g., oxygen and glucose). To examine the link between microvessel structure and its influence on blood perfusion in the brain, we applied a mathematical approach to infer degree of vascular blood flow (permeability) and directionality of the microvascular network (Figure 3a; Movie S5; Table S6). To limit our simulation to small vessels, we removed large vessels with radius > 4 μm. The permeability was calculated by solving the Hagen–Poiseuille equation system of the entire network in the control volume, taking into account vascular density, connectivity, and network topology (e.g., tortuosity) (Figure 3a-b). Thus, our permeability measurement can provide estimates of blood perfusion at a given pressure.

**Figure 3.**
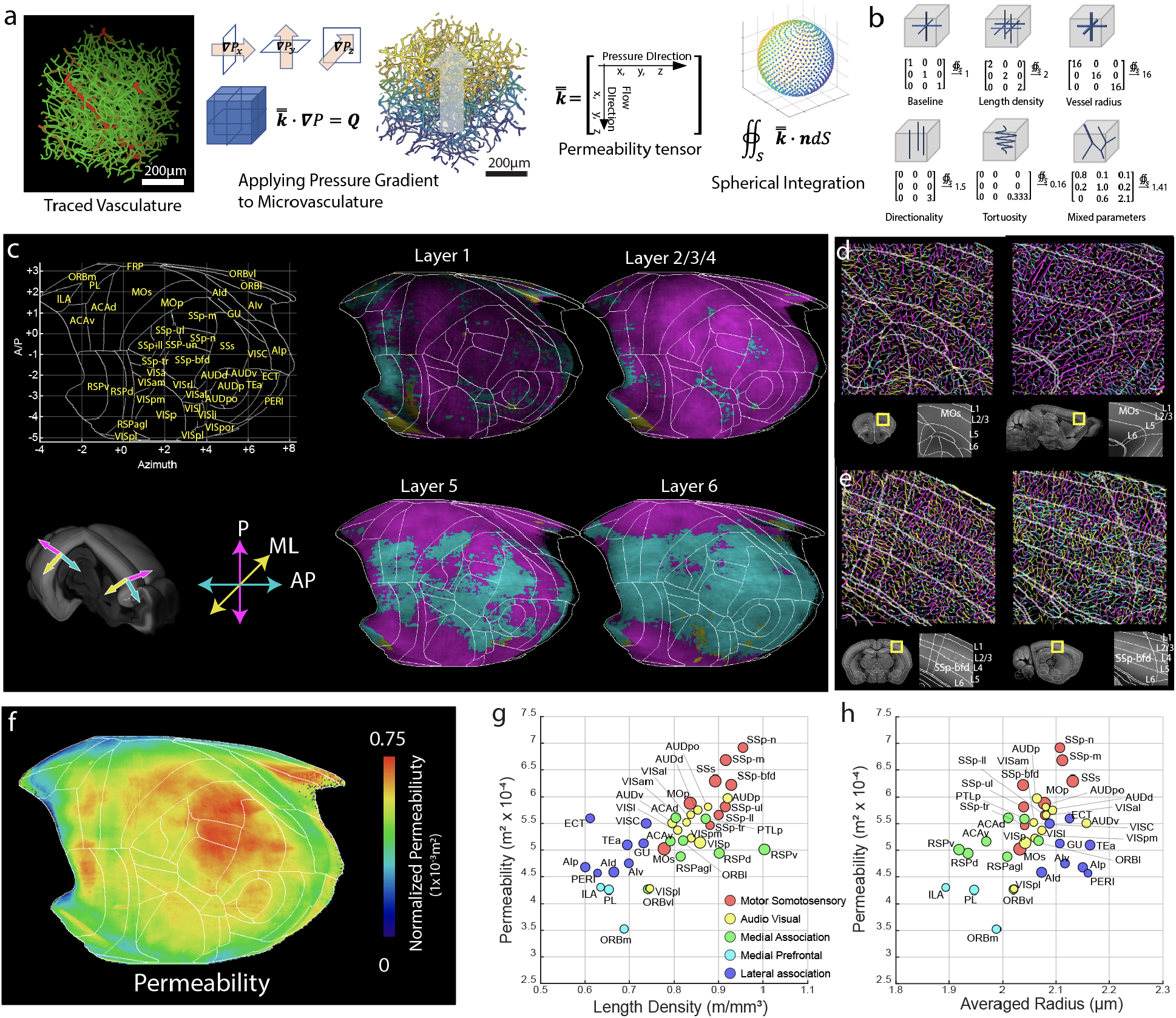
Anisotropy of the cerebral microvascular network and its impact on permeability. **a**. The flow chart of the permeability fluid dynamic simulation. From left to right: Original traced data (red for large vessel with radius > 5 μm and green for small vessels), applying pressure profile on the surface of the control volume with a gradient profile and solving the coincide flux equation set for the permeability tensor, the annotation rule of the permeability tensor, and the sampling dots of the numerical spherical integration. Equation symbols: *l*: perfusing length, *P*: pressure, *Q*: Blood flux, *R*: resistance, *μ*: viscosity, *k*: permeability, Δ: changing of the quantity, ∙: dot product, ∇: gradient, ∯ *s*: spherical integral, *S*: spherical surface, *n*: normal direction, bold-font: vector, double-top-bar: tensor. **b**. Examples illustrating how the structure of the vasculature network impacts the permeability tensor. **c-e**. Microvessel anisotropy measurement in the isocortex. **c**. Microvessel directionality in cortical layers. Only the dominant direction is displayed for simplicity. Microvessels are colored based on its directionality; magenta for penetrating (P), cyan for anterior-posterior (AP), yellow for medial-lateral (ML). Directionality axis is based on cortical column angles in each area. Examples from the motor cortex (**d**) and somatosensory (**e**) cortices. **f**. Permeability results in the cortical flatmap. **g-h**. Relationship between permeability and vessel length density (**g**), or average radius (**h**). Please see Table S6 for full data and abbreviation.

We first examined how the geometry of microvascular networks can influence directionality of blood flow by calculating the microvessel anisotropy using the permeability tensor in the isocortex (Figure 3c). We used three axes according to the cortical column direction: penetrating (P) axis along the cortical column as a main blood input direction from surface vessels, anterior-posterior (AP) and medial-lateral (ML) as two vascular communication directions within areas (Figure 3c). Our analysis showed that microvessels oriented in the P-axis dominated in the anterior (e.g., prefrontal and motor area) and posterior cortical areas (e.g., visual area) while mid-cortical areas (e.g., somatosensory area) showed vasculature orientation in the P and AP axes dominating the superficial layer and the deep layers, respectively (Figure 3c). For instance, the secondary motor cortex (MOs) shows dominant P-axis vasculature (magenta) while the SSp-bfd shows a clear switch to AP axis (cyan) vasculature preferentially between layers 4 – 6 (Figure 3d-e). This result suggests that mid-cortical areas containing many primary motor-sensory areas have a high degree of anterior-posterior vascular communication in deep layers to facilitate blood perfusion across these hyper-vascularized areas.

Next, we applied the permeability measurement across the isocortex and plotted the result in the cortical flatmap (Figure 3f). Overall, motor sensory groups showed higher permeability than association groups in correlation with the vascular density (Figure 2g-h; Table S6). Particularly, SSp-nose (n), mouth (m), and barrel field (bfd) are areas with the highest permeability in the isocortex (Figure 3f-h). Noticeable exceptions include the RSPv with relatively low permeability despite highest cortical vessel density due to small vessel radius, and a few lateral association areas (e.g., the ectorhinal cortex; ECT) which has relatively high permeability due to large vessel radius (Figure 3g-h).

Collectively, our data suggests that microvessels in primary motor sensory cortices are structured to provide high degree of blood perfusion compared to other cortical areas.

### Pericyte and nNOS neuron density mapping reveals differential vasoregulation between cortical areas

Pericytes, a mural cell type, are proposed to actively regulate microvascular diameter and permeability ^6,8,38–40^. Here, we ask whether pericyte distribution shows distinct densities across brain areas in support of distinct local brain functions, in a similar fashion to cortical vasculature. We found that pericyte density across cortical areas showed a similar distribution pattern to the vascular density (Figure 4a-c; Tables S3; Movie S2). Overall, primary sensory areas as well as the RSPv showed higher pericyte densities than association (medial, lateral association and medial prefrontal) groups (Figure 4a-c). A very strong positive correlation between pericyte and vascular densities suggests that the number of pericytes per length of vasculature is overall constant across different cortical areas (Figure 4c). However, when we examined layer-specific differences in the density of pericytes and vasculature, we observed that relative pericyte density as well as the ratio between pericyte and vascular densities were highest in layer 5 (Figure 4e; Table S3). This suggests that layer 5, characterized by its large pyramidal neurons, may require higher pericyte coverage per vasculature to finely tune the regulation of blood flow compared to other cortical layers.

**Figure 4:**
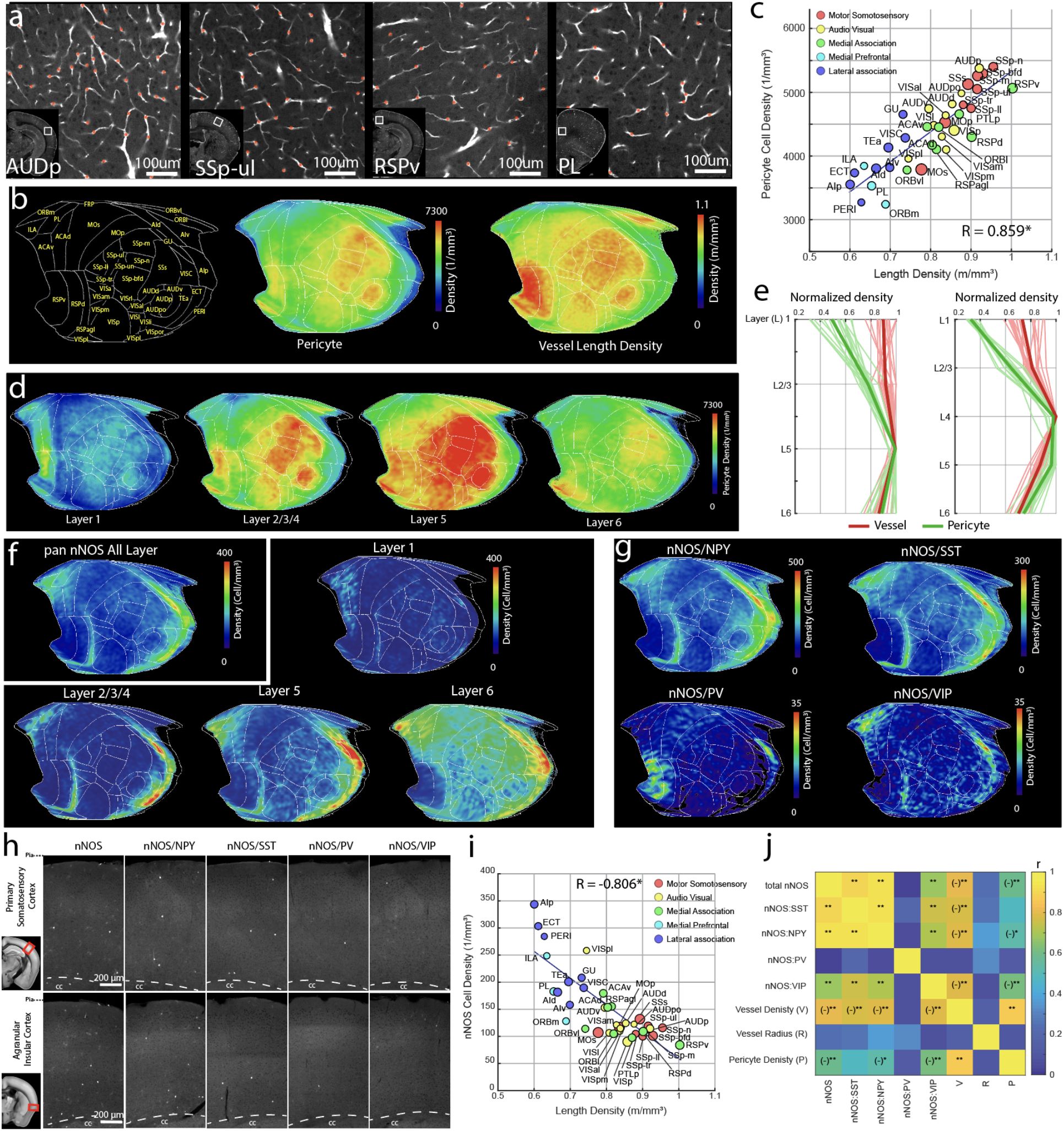
Cortical pericyte and nNOS neuron densities display opposite correlation with vascular density. **a.** Example images of areas showing variability in pericyte density from PDGFRβ-Cre:Ai14 with pericyte detection algorithm (red stars). **b.** Cortical flatmap of averaged pericyte density across the isocortex in comparison to the vessel length density. **c.** Scatter plot demonstrating significantly positive correlation between pericyte density and vascular length density in isocortical regions (R=0.859, p value =1.86×10^−12^). See Table S3 for abbreviation. **d**. layer specific pericyte distribution. **e**. relative pericyte (green) and vessel (red) density (normalized against maximum value within the area) in cortical regions without (left) or with the layer 4 (right). **f-g.** Averaged density of pan nNOS neurons (**f**) or nNOS subtypes (**g**) on the cortical flatmap. Note higher nNOS density in the medial prefrontal and lateral association areas. **h.** Representative STPT images of nNOS cell types from the primary somatosensory cortex and the agranular insular cortex. **i.** Scatter plot showing significant negative correlation between total nNOS density and vessel length density in the isocortex (R value =-0.806, p value = 6.9 × 10^−7^). **j.** Correlation matrix between nNOS cell types and vascular/pericyte measurements. *p<0.05, **p<0.005 after the Bonferroni correction. (-) denote negative correlation.

Next, we examined the relationship between the cerebrovascular network with pericytes and the distribution of cortical nNOS-expressing neurons whose activity was shown to cause vasodilation ^22–24^. Based on our results so far, our expectation was that the density distribution of nNOS neurons may follow the distinct patterns of vasculature and pericyte densities in providing a more robust blood flow support for the motor-sensory versus prefrontal-association cortical areas. Unexpectedly, while we did observe up to two-fold differences in nNOS neuronal density across isocortical regions (Figure 4f; Table S4; Movie S3), nNOS neuron density was higher in the association cortical areas compared to the motor-sensory areas, including the RSPv – the opposite pattern than that of the vascular and pericyte densities (Figure 4f,h-j). We also noted that the highest density of nNOS neurons was found in layer 6 in all cortical areas compared to the highest densities of vasculature and pericytes in layer 5 (Figure 4f). For nNOS subtypes, nNOS/NPY, nNOS/SST, and nNOS/VIP subtypes showed similar density patterns as the pan nNOS neurons (Figure 4g). In contrast, nNOS/PV neurons, despite having much lower density, showed relatively higher expression in the RSPv (Figure 4g), suggesting a subtype-specific role in this cortical area. The contrasting nNOS distribution is further evident from correlation analysis for the relationship between nNOS neuronal densities and vascular density across different areas, which revealed a significant negative correlation in the isocortex that are true for all subtypes except nNOS/PV (Figure 4i-j). Similarly, nNOS neurons, including nNOS/NPY and nNOS/VIP subtypes, showed significant negative correlation with pericyte density (Figure 4j). In contrast, nNOS neurons and all of their subtypes did not show any correlation with average vasculature radius (Figure 4j). This data shows a surprisingly distinct distribution of nNOS neurons across cortical areas with overall stronger nNOS-based vasomotor regulation in association cortices than in primary motor sensory cortices.

### Cortical PV interneurons and glutamatergic neurons show positive correlation with the vascular network

Glutamatergic and GABAergic neuronal cell types have different energy consumption and metabolic costs ^41^. We examined whether glutamatergic neurons and specific GABAergic neuronal subtypes show any significant correlation with vascular and pericyte distribution in the isocortex (Figure 5; Table S5; Movie S4). Density plotting using our isocortical flatmap allowed us to visualize distinct neuronal cell type distributions and localizations across the isocortex, revealing a clear pattern when compared to the vessel length and pericyte densities (Figure 5a). First, pan-glutamatergic neurons (vGlut1+), but not pan GABAergic (Gad2+) neurons, showed modest, but significant positive correlation with vascular and pericyte density (Figure 5a-d). However, among different GABAergic cell types, PV+ interneurons showed a strikingly strong positive correlation with the vascular length density whereas the other interneuron subtypes (SST+ and VIP+) did not show a significant correlation with vascular density (Figure 5c). All motor sensory groups and the RSPv showed high PV+ interneuron density comparable to the vascular density (Figure 5a,c). As expected, the pericyte distribution followed the similar correlation patterns as the vessel length density with all of the neuronal subtypes studied (Figure 5d). Importantly, cortical PV neurons are involved in the generation of gamma-band oscillations ^42–44^, which are linked with increased vasodilation and blood flow in the brain ^45^. Thus, our results suggest that correlated vascular, pericyte, and PV+ interneuron densities may act to support local gamma-band oscillation neural activity during rapid signal processing in sensory cortices.

**Figure 5:**
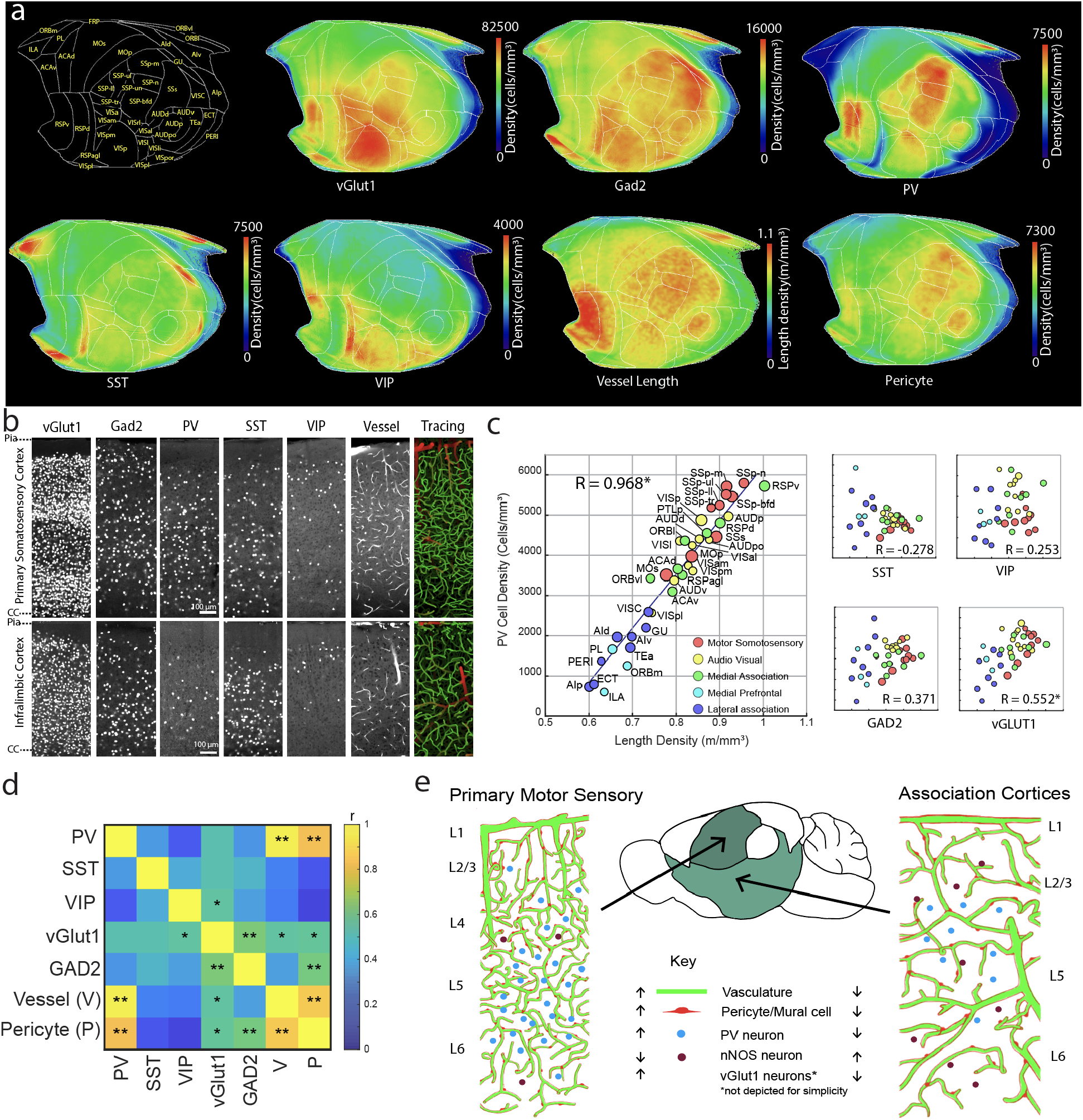
Cortical parvalbumin+ and vGlut1+ neurons positively correlated with vasculature density. **a.** Cortical flatmap showing density distributions of neuronal subtypes as well as vessel length and pericyte densities. **b**. Examples of neuronal cell types, and the vasculature and its tracing result (large vessels=red, microvasculature=green) from the densely vascularized primary somatosensory and sparsely vascularized infralimbic cortices. **c.** Correlation between vascular density and neuronal subtypes. Note very strong positive correlation with PV density (R = 0.968, p value = 8.5×10^−22^) and positive correlation with vGlut1 excitatory neuronal density (R= 0.552, p value = 5.9×10^−3^). **d**. Correlation matrix between neuronal subtypes, vessel length density and pericyte density. *p<0.05, **p<0.005 after the Bonferroni correction. **e.** Cortical organization of the vascular/pericyte network and neuronal cell types. Primary motor sensory cortices are characterized by relatively high density of vessels, pericytes, PV interneurons, and vGlut1 excitatory neurons, and a low density of nNOS neurons. In contrast, association cortices show the opposite pattern.

In summary, our data suggest that the isocortex in mice is composed of two domains with distinct vascular/pericyte and neuronal cell types composition; 1) the primary motor-sensory domain (motor, somatosensory, audio, visual cortices, and the RSPv) with a high density of vessels and pericytes positively correlated to the density of PV+ inhibitory neurons and less prominently vGlut2+ excitatory neurons, but negatively correlated to the density of nNOS neurons, 2) the association domain (lateral, medial, and medial prefrontal) which comprise the opposite pattern of vasculature, pericytes, and neuronal density distribution (Figure 5e; Table 2).

**Table 2:** Density of the cerebrovasculature, pericytes, and neuronal subtypes in the isocortex. Number = average ± standard deviation, see Table S1, S3, S4, S5 for full dataset.

| Region of Interest | Vessel length density (m/mm<sup>3</sup>) | Pericyte (cell/mm<sup>3</sup>) | nNOS neurons (cell/mm<sup>3</sup>) | PV neurons (cell/mm<sup>3</sup>) | vGlut neurons (cell/mm<sup>3</sup>) |
| --- | --- | --- | --- | --- | --- |
| Motor | 0.81 ± 0.05 | 4,164 ± 308 | 104 ± 17 | 2,678 ± 203 | 35,566 ± 823 |
| Somatosensory | 0.91 ± 0.07 | 5,177 ± 466 | 108 ± 16 | 3,664 ± 281 | 45,022 ± 838 |
| Auditory | 0.86 ± 0.06 | 5,049 ± 535 | 121 ± 16 | 2,965 ± 295 | 42,178 ± 1,676 |
| Visual | 0.82 ± 0.07 | 4,071 ± 440 | 112 ± 22 | 2,927 ± 352 | 49,329 ± 1,132 |
| Retrosplenial | 0.92 ± 0.07 | 4,598 ± 548 | 103 ± 20 | 3,330 ± 387 | 44,631 ± 2,494 |
| Posterior Parietal | 0.87 ± 0.07 | 4,684 ± 404 | 91 ± 17 | 3,220 ± 255 | 47,030 ± 875 |
| Orbital | 0.75 ± 0.08 | 4,685 ± 404 | 106 ± 17 | 2,290 ± 320 | 38,014 ± 1,559 |
| Anterior Cingulate | 0.8 ± 0.05 | 4,324 ± 350 | 154 ± 32 | 2,371 ± 213 | 41,208 ± 1,039 |
| Prelimbic | 0.65 ± 0.04 | 3,560 ± 458 | 171 ± 34 | 1,199 ± 171 | 39,270 ± 1,098 |
| Infralimbic | 0.64 ± 0.02 | 3,872 ± 380 | 232 ± 38 | 503 ± 120 | 43,679 ± 1,973 |
| Visceral | 0.74 ± 0.04 | 4,312 ± 434 | 177 ± 33 | 1,846 ± 204 | 37,081 ± 1,546 |
| Gustatory | 0.73 ± 0.03 | 4,681 ± 366 | 194 ± 34 | 1,652 ± 212 | 40,783 ± 1,467 |
| Agranular Insular | 0.65 ± 0.05 | 3,763 ± 315 | 207 ± 41 | 1,154 ± 118 | 31,290 ± 1,023 |
| Perirhinal | 0.63 ± 0.03 | 3,300 ± 388 | 265 ± 69 | 979 ± 142 | 24,789 ± 3,196 |
| Temporal Association | 0.7 ± 0.04 | 4,160 ± 403 | 187 ± 37 | 1,189 ± 210 | 42,407 ± 1,907 |
| Ectorhinal | 0.61 ± 0.02 | 3,763 ± 321 | 283 ± 63 | 623 ± 117 | 38,578 ± 3,400 |

### Sensory thalamo-cortico-striatal circuit shares high densities of vasculature and pericyte

Next, we asked whether the high vascular and pericyte density specific to primary motor-sensory vs association cortices is also shared across thalamo-cortico-striatal pathways. We used thalamo-cortical and cortico-striatal connectivity datasets to identify clusters of thalamic and dorsal striatal areas that are well connected with the five cortical domains ^46–49^. In the thalamus, sensory thalamic areas processing somatosensory (e.g., ventral posteromedial thalamus; VPM), auditory (e.g., medial geniculate complex; MG), and visual (e.g., dorsal lateral geniculate complex; LGd) information as well as the anteroventral nucleus of thalamus (AV), which is strongly connected with the RSPv, showed higher densities of vasculature and pericytes than non-sensory thalamic areas connected to medial prefrontal and other association cortical groups (e.g., nucleus of reuniens; RE) (Figure 6a-b).

**Figure 6:**
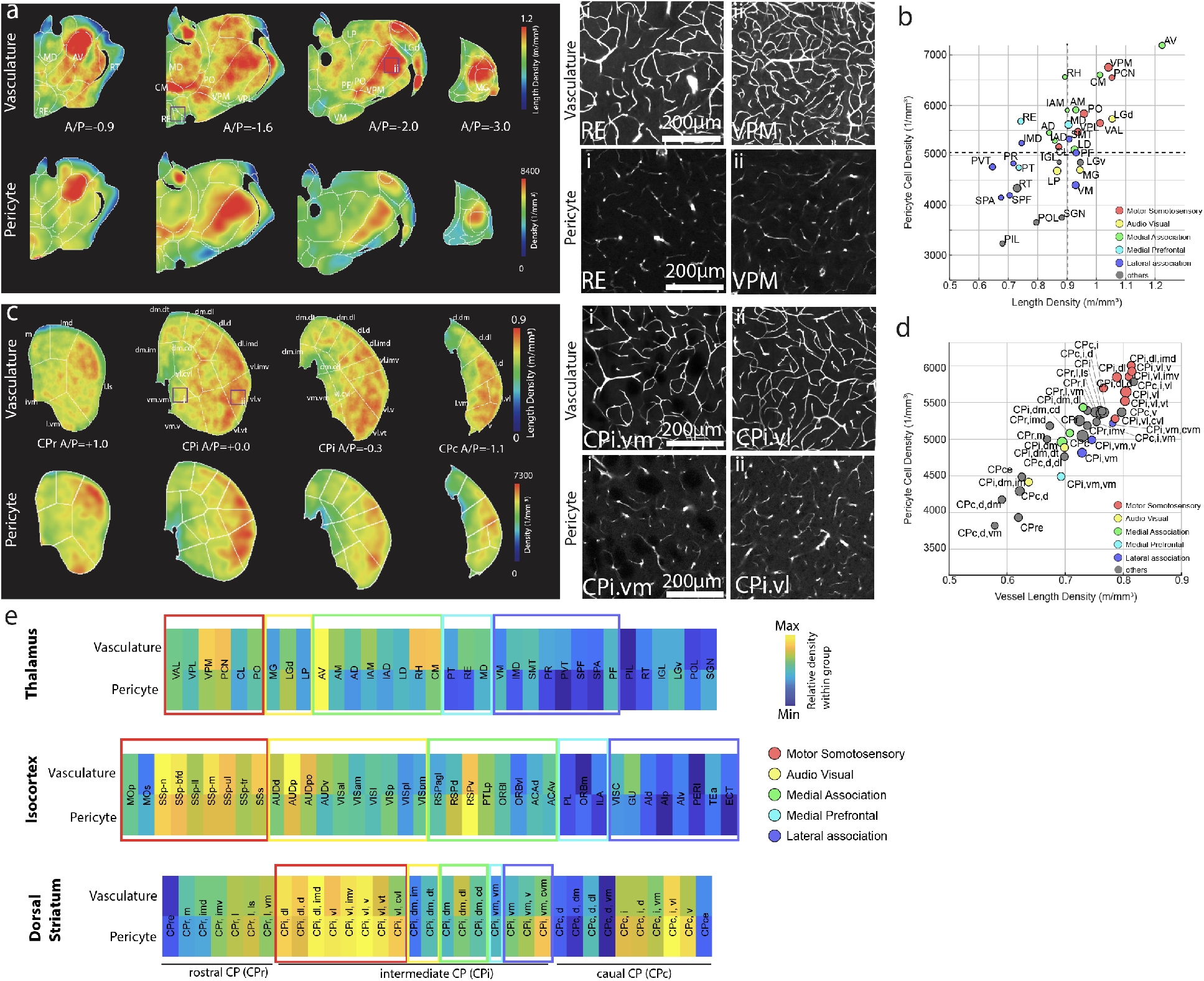
high density of vascular/pericyte network in motor sensory thalamo-cortico-striatal pathway. **a**. Heatmap of vascular (top) and pericyte (bottom) densities in the thalamus (left side) with examples from nucleus of reuniens (RE, low densities) and the ventral posteromedial thalamus (VPM, high densities) in the right side. **b**. Density scatter plot of pericytes and the vessel length densities. Colors of thalamic areas are assigned based on anatomical connectivity with specific cortical groups. **c**. Heatmap of vascular (top) and pericyte (bottom) densities in the striatum (left side) with examples from the intermediate CP ventral medial (CPi.vm, low densities) and the intermediate CP ventral lateral (CPi.vl, high densities) in the right side. **d**. Density scatter plot of pericytes and the vessel length densities. Colors of striatal areas are the same as (**b**). **e**. Heatmap of vascular and pericyte densities normalized within anatomical areas. Color of boxes represents cortical groups and their connected thalamo-cortical areas. Note that motor sensory areas contain higher densities compared to association areas. See Table S1 and S7 for abbreviations.

We then examined vascular and pericyte densities in subregions of the dorsal striatum (caudate putamen; CP) using detailed anatomical segmentations established in the Allen CCF ^50^. The intermediate CP (CPi) receives topographically segregated projections from cortical domains while rostral (CPr) and caudal (CPc) areas have inter-mixed projections from many cortical domains ^46,48^. Our analysis revealed that CPi areas receiving input from primary motor-sensory cortices such as the CPi ventral lateral (CPi.vl) had relatively higher vascular and pericyte densities than the CPi ventral medial (CPi.vm) areas connected with medial prefrontal and lateral association cortical projections (Figure 6c-d; Table S7). Heatmap plots of relative vascular and pericyte densities with each anatomical region showed the clear pattern of primary motor-sensory processing areas comprising overall higher vascular and pericyte densities compared to association areas throughout the thalamo-cortico-striatal pathways (Figure 6e).

### Data resources to further examine brain energy supply and regulation

While we focused our analysis primarily on the isocortex, our brain-wide high resolution vasculature and cell type mappings open new opportunities to understand global and local energy supply and its regulation. For instance, we provide our brain-wide permeability simulation result and its relationship with pericyte density and pericyte coverage (pericyte per vessel length density) as a resource to elucidate regional blood perfusion and vascular integrity (Figure S4). We noted that many hippocampal areas including the dentate gyrus (DG) and the subiculum (SUB) showed low permeability, pericyte density, and pericyte coverages compared to other cortical and subcortical areas, which may confer regional vulnerabilities upon pathological conditions ^51,52^ (Figure S4; Table S6; Movie S5).

We also include a freely downloadable simulation ready dataset to model vascular flow in the whole brain as well as high resolution raw images in a publicly available database. To facilitate ease of access and intuitive visualization to examine large scale imaging datasets, we created a web-based resource (https://kimlab.io/brain-map/nvu/) that displays navigable z-stacks of full-resolution images for our STPT datasets including FITC filled vasculature, PDGFRβ-Cre:Ai14 for pericytes, nNOS-Cre:Ai14 for total nNOS neurons, and nNOS-Cre:Ai65:(NPY, SST, PV, or VIP-Flp) for nNOS subtypes. This web-based resource also provides interactive 3D visualizations, allowing users to navigate our quantitative vascular and cell type measurements registered in the Allen CCF.

One notable new cell type resource is the nNOS neuronal subtype brain-wide distribution data. In addition to cortical nNOS neurons, nNOS neurons in subcortical areas including the cerebellum also powerfully regulate neurovascular coupling ^53,54^. Our results indicate that the total nNOS neuronal density is highest in the accessory olfactory bulb (AOB), followed by the cerebellum, the medial amygdala (MEA), the dorsal medial hypothalamus (DMH) (Figure 7a-c; Table S4). In contrast, the isocortex, the hippocampus, and the thalamus showed overall low nNOS neuronal density (Figure 7c). Of nNOS subtypes, the nNOS/NPY neurons represent the majority of nNOS subtypes in cerebral cortical, hippocampal, and cerebral nuclei areas (Figure 7a-c). The nNOS/SST subtype showed overall similar density compared to the nNOS/NPY subtype except in hippocampal regions which had noticeably low density (Figure 7a-c). The nNOS/PV subtype showed overall low density across the whole brain, except for very high density in the cerebellum (Figure 7a-c). Lastly, the nNOS/VIP subtype had the lowest density compared to the other nNOS^+^ subtypes with sparse expression in a few areas such as the subiculum of the hippocampus (SUB) (Figure 7a-c). Noticeably, many amygdala and hypothalamic areas as well as the AOB showed high nNOS density that was not reflected in the nNOS interneuron subtype populations, suggesting that nNOS neurons in these areas may represent different nNOS subtypes ^55^. Thus, the current brain-wide nNOS subtype mapping unveils region specific distributions of the vasomotor neurons.

**Figure 7.**
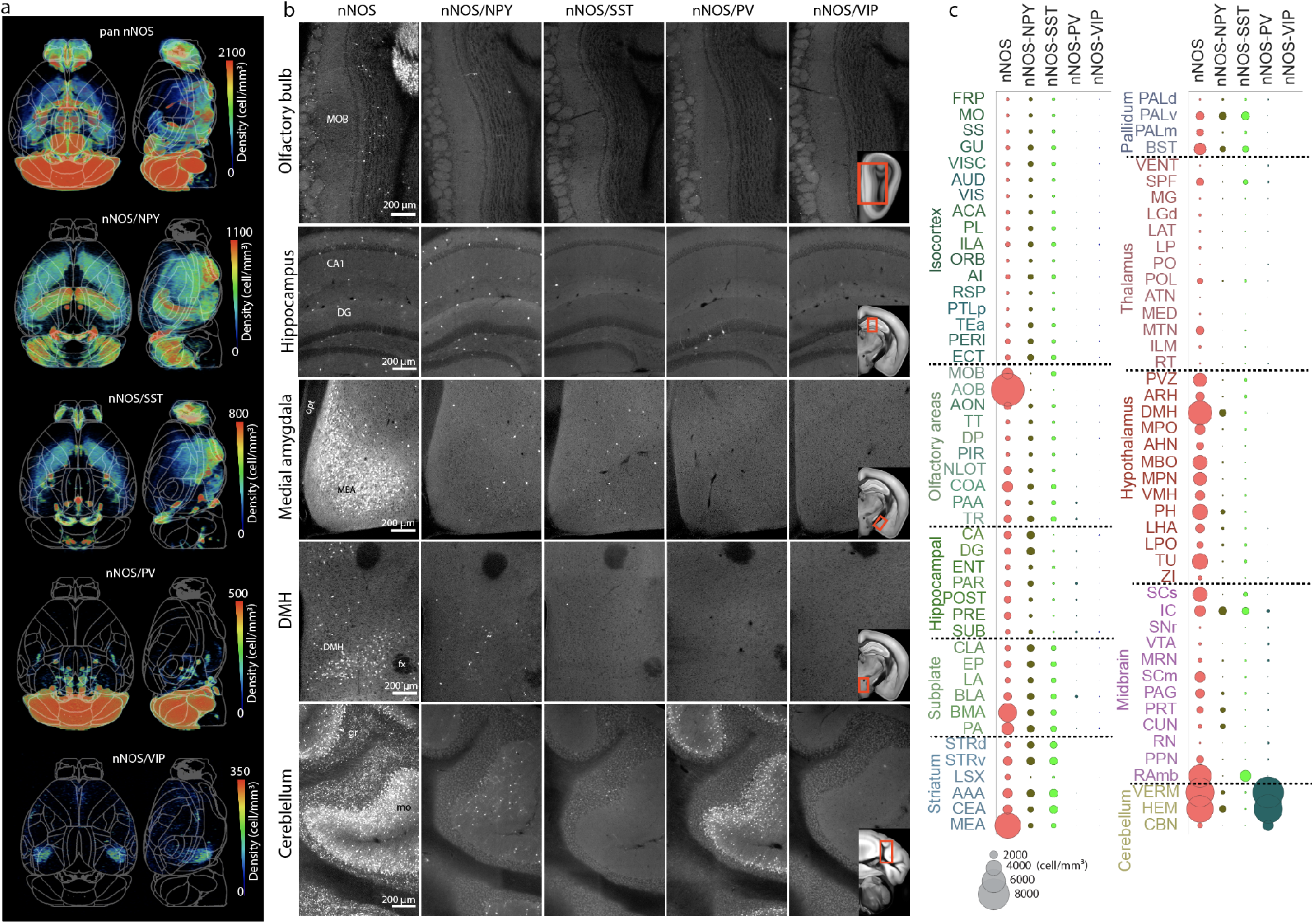
Brain-wide density map of nNOS neurons and their subtypes. **a.** Heat maps demonstrating the distribution of total nNOS and nNOS subtype populations. See also Movie S4 and Table S4. **b.** Representative raw images of nNOS, nNOS/NPY, nNOS/SST, nNOS/PV and nNOS/VIP neurons in the olfactory bulb, hippocampus, medial amygdala (MEA), dorsomedial hypothalamus (DMH) and cerebellum. Reference atlas images included on nNOS/VIP images show the area displayed for each region of interest. Main olfactory bulb (MOB), Ammon’s horn (CA1), dentate gyrus (DG), optic tract (opt), fornix (fx), granular (gr), molecular (mo). **c.** Graph showing nNOS density by brain region for the total nNOS neurons and their subtypes. Size of circle corresponds to density as shown in the key at the bottom. Please see Table S4 for full names of abbreviations.

## Discussion

The structural organization of regional vascular networks is crucial to support local brain function and may reflect susceptibility to different pathologies. Here we present cellular resolution maps of cerebral vasculature, pericytes, and neuronal subtypes in the mouse brain. Our cerebrovascular map, in combination with fluid dynamics simulations, reveals organizational principles of microvessels and pericytes in relationships to several key neuronal cell types, highlighting heterogenous blood perfusion and potential differences in blood flow regulation across different brain regions. Most notably, we observed striking correlations between vascular density and parvalbumin interneuron and vasomotor nNOS neuron densities in the isocortex, suggesting that regional differences in neuronal cell type composition are linked with different energy demands and are met by region-specific patterns of cortical vascularization. In combination with web visualization and downloadable datasets, our maps serve as a comprehensive resource to examine microvasculature and associated cell types.

### Cortical neuronal cell types and the vascular network

A prevailing theory of cortical organization is that the cortex is composed of repeating cortical columns with a common microcircuit motif ^56^. For example, excitatory and inhibitory neuronal subtypes make stereotypic local connection patterns, called canonical cortical circuits, that are similarly observed across cortical areas ^57,58^. However, this view has been challenged by recent data that different cortical domains show distinct cell type compositions and hemodynamic responses ^15,20^. Results from the current study provide further evidence that vascular networks, including pericytes and vasomotor neurons, are organized in distinct spatial patterns to meet energy demands from motor-sensory and association cortices ^3,16^.

Motor and sensory signals require precise temporal and spatial information processing in primary motor sensory cortices to perceive dynamic external signals and to execute motor commands. Although previously included as a part of the association cortices, we consider the RSPv as a part of the sensory area because of its role in processing rapid navigational information from the dorsal subiculum ^37^. In contrast, association cortices integrate information from broader areas with slower temporal kinetics. We previously identified a higher density of PV neurons in sensory cortices compared to association cortices ^20^. Cortical PV neurons are fast spiking interneurons that participate in generating gamma oscillations and are one of the most energy demanding neurons ^21,42,44,59,60^. Thus, our current results suggest that a high density of microvessels and capillary pericytes in the sensory cortices provide an efficient energy support system for PV dominated local circuits to accommodate high energy consumption and to mediate local functional hyperemia for rapid sensory processing. In contrast, association cortices contain relatively high densities of nNOS neurons despite low vascular, pericyte, and PV densities. Although nNOS interneurons represent only about 2% of cortical neurons, activation of nNOS neurons robustly dilates cerebral arterioles to generate increases in cerebral blood flow ^22–24^. Thus, the relatively higher density of nNOS neurons in association areas suggests that this cell type can exert more powerful vasodilation in larger areas to compensate for a lower vascular density in these high cognitive areas.

Moreover, we also found that areas of the thalamus and the dorsal striatum heavily connected to motor-sensory cortices contain high densities of vasculature and pericytes compared to thalamo-striatal areas linked with association cortices. Thus, our data demonstrate that this dense network of vasculature and pericytes is conserved throughout neural circuits processing primary motor-sensory information.

### Comprehensive data resources to understand relationships between brain regional vascular organization and energy homeostasis

Although recent approaches using light sheet microscopy to examine fine cerebrovascular structure have provided advantages in rapid data acquisition as well as 3D immunolabeling to mark different vascular compartments ^26,27^, the required tissue clearing methods can introduce microscopic volume distortions, which can lead to inconsistent measurements ^25^. Here, we used serial two-photon tomography to visualize the whole cerebrovasculature at single capillary resolution from intact mouse brains, revealing a cerebrovascular map that closely represents physiological conditions as confirmed by *in vivo* two-photon microscopy. Thus, our dataset allows us to perform precise computational simulations to estimate blood flow efficiency based on structural arrangements of microvessels, including deep cortical layers as well as subcortical areas, which are hard to access with *in vivo* two-photon microscopy. For example, our simulation result showed that cerebrovasculature in primary motor sensory cortices is structured to provide higher blood perfusion with stronger anterior-posterior vascular connection in the deep layer than association cortices. Future works using our data resources, including computational modeling considering additional information (e.g., blood pressure and viscosity), can help to gain a more complete understanding of brain blood perfusion and its change with additional risk factors such as strokes ^61,62^. Moreover, our high resolution cerebrovascular maps can provide a detailed structural basis of signals for functional neuroimaging modalities such as functional magnetic resonance imaging or newly emerging functional ultrasound imaging ^63^

Our study also presents the first brain-wide quantitative pericyte map. Previous functional studies identified that pericytes actively regulate the diameter and permeability of microvessels ^6,8^. Our results complement previous work by providing capillary pericyte population density across the brain. We observed a strong positive relationship between pericyte and vascular density in the cortex, suggesting that pericyte coverage per microvessel remains similar across different cortical areas in the normal adult mouse brain. Notably, we found the highest pericyte coverage per vascular length in layer 5 across all cortices. Since large pyramidal neurons in layer 5 act as main cortical output to the rest of the brain, the high density of pericytes may confer extra control over blood flow in this energy demanding layer. Moreover, our subcortical mapping results provide new opportunities to investigate pericyte arrangements in largely under-studied brain regions. For instance, thalamic areas have overall higher pericyte density compared to other areas (Figure S4). Interestingly, thalamic pericytes have shown resistance to disrupted PDGFRβ signaling, while cortical and striatal pericytes were more vulnerable ^64^. The combination of high density and cellular resilience may confer extra protection to maintain vascular integrity in the thalamus. Conversely, relatively low pericyte density in the hippocampal areas and association cortices can make these areas more vulnerable to pathological conditions^4,65,66^.

Lastly, our nNOS results with added subtype specificity offer new insight to understand nNOS subtype coverage across the whole brain. In contrast to well-studied cortical nNOS neurons, the function and vasomotor characteristics of subcortical nNOS neurons is largely unknown. Previous studies suggested that nNOS signaling in the cerebellum and the hypothalamus is linked with neurovascular coupling ^53,54^. Our comprehensive nNOS and nNOS subtype maps can guide future research to determine what brain regions and which nNOS subtypes need to be examined to establish a causal relationship between nNOS neuronal types and local hemodynamic response.

In summary, our quantitative information on cerebrovasculature and associated cell types establishes a platform for future studies to gain a deeper understanding of how energy demand and supply maintain balance in a normal brain from a cellular architectural perspective and how this homeostatic mechanism changes under pathological conditions.

## Material and Methods

### Animals

Animal experiments were approved by the Institutional Animal Care and Use Committee at Penn State University and Cold Spring Harbor Laboratory. For all genotypes in this study, both adult male and female mice were used. Adult 2-month-old C57BL/6 mice were bred from C57BL/6 mice directly obtained from the Jackson Laboratory and used for vascular tracing experiments with FITC filling (n=4). For pericyte specific experiments, male PDGFRβ-Cre mice ^32^ were crossed with female Ai14 mice (Jax: Stock No: 007914) as previously described ^1^. These PDGFRβ-Cre:Ai14 mice exhibit PDGFRβ-driven tdTomato expression in two distinct vascular cell types, pericytes and vascular smooth muscle cells (vSMCs). For isocortical cell types, vGlut1-Cre (Jax: 023527) and Gad2-Cre (Jax: 010802) mice were crossed with Ai75 reporter mice (Jax: 025106). nNOS-CreER mice were used to label nNOS neurons (Jax: Stock No: 014541)^33^. After nNOS-CreER mice were crossed with Ai14 mice, the nNOS-CreER:Ai14 offspring were administered with an intraperitoneal (i.p.) tamoxifen (Sigma, cat.no. T5648-1G) injection (100mg/kg) at P16. Similarly, for nNOS-subtypes, nNOS-CreER mice were initially crossed with Ai65 mice (Jax; Stock No: 021875), which were further crossed with PV-flp (Jax Stock No: 022730), SST-flp (Jax Stock No: 028579), NPY-flp (Jax Stock No: 030211), or VIP-flp (Jax Stock No: 028578) mouse lines, to generate triple transgenic mice which allowed for tdTomato fluorescent labeling of nNOS expression within these interneuron populations. To allow for postnatal specific expression of tdTomato in nNOS+ subtype populations, tamoxifen injections dosed at 75mg/kg were given at P10, P12, and P14 timepoints. We used 10 animals for each PDGFRβ:Ai14, nNOS:Ai14, nNOS:VIP:Ai65, 9 animals for nNOS:NPY:Ai65 and nNOS:PV:Ai65, 8 animals for nNOS:SST:Ai65 from both males and females as well as 7 animals (all males) for vGlut1:Ai75, and 9 animals (all males) for Gad2:Ai75. All animals were used once to generate data. We used tail genomic DNA with PCR for genotyping. Brain samples were collected at 2 months old age for all mouse lines.

### Perfusion and tissue processing for STPT imaging

Animals were deeply anesthetized with a ketamine-xylazine mixture (100 mg/kg ketamine, 10 mg/kg xylazine, i.p. injection) for both regular perfusion and vascular labeling. Transcardiac perfusion with a peristaltic pump (Ismatec, cat.no.: EW-78018-02) was used with 1X PBS followed by 4% paraformaldehyde, both injected through a small incision in the left ventricle, in order to wash out blood and allow for tissue fixation, respectively. Brains were dissected carefully in order to preserve all structures. For vessel labeling, transcardiac perfusion with a peristaltic pump (Welch, Model 3100) was used with 1X PBS followed by 4% paraformaldehyde at 0.3 ml/min, in order to wash out blood and for tissue fixation, respectively. To ensure that the large surface vessels would remain filled with the gel perfusate, the body of the mouse was tilted by 30° before gel perfusion (with the head tilted down), as previously described ^30^. Following the fixative perfusion, the mouse was perfused at 0.6 ml/min with 5 ml of a 0.1% (w/v) fluorescein isothiocyanate (FITC) conjugated albumin (Sigma-Aldrich, cat.no.: A9771-1G) in a 2% (w/v) solution of porcine skin gelatin (Sigma-Aldrich, cat.no: G1890-500G) in 1X PBS. Immediately after perfusion, the heart, ascending and descending aorta as well as the superior vena cava, were all clamped with a hemostat (while the butterfly needle was simultaneously removed from the left ventricle). This served to prevent any pressure changes in or gel leakage from the brain vasculature. Next, the entire mouse body was submerged in an ice bath to rapidly solidify the gel in the vessels. Then, the head was fixed in 4% PFA for one week, followed by careful dissection of the brain to avoid damage to pial vessels. After fixation and dissection, the brain was placed in 0.05M PB until imaging. Any animals that had poor perfusion and/or possible air bubbles interfering with the gel perfusion were excluded from imaging and any further analysis.

### Serial two photon tomography (STPT) imaging

Prior to imaging, the brain sample was embedded in oxidized agarose and cross-linked in 0.05M sodium borohydrate at 4ºC for at least 2 days ahead of imaging ^20,67^. This procedure allows for seamless cutting of 50μm thick sections, while also preventing any tearing of the brain surface. The embedded brain sample was then glued to the sample holder and fully submerged in 0.05M PB in an imaging chamber. For STPT imaging (TissueCyte), we used 910nm excitation using a femtosecond laser (Coherent Ultra II) for all samples. Signals in the green and red spectrum were simultaneously collected using a 560 nm dichroic mirror (Chroma). For pericyte and neuronal subtypes, STPT imaging was conducted with 1×1 μm (*x,y*) resolution in every 50 μm (*z*), with the imaging plane set at 40μm deep from the surface, as previously described ^20,67^. For vascular imaging, optical imaging (5 μm z step, 10 steps to cover 50μm in *z*) was added in the imaging, producing 1×1×5 μm (*x,y,z*) resolution beginning at 20μm deep from the surface. Due to length of imaging time required for vascular imaging, each brain sample was imaged through multiple imaging runs to adjust the imaging window size in order to reduce overall imaging time.

### Computational: STPT Image reconstruction

To measure and correct an optical aberration from the objective lens (Figure S1), we imaged a 25 μm EM-grid (SPI supplies, cat.no.: 2145C) as a ground truth for spatial data ^68^. We annotated all cross points of the grid and computed the B-spline transformation profile from the grid image to the orthogonal coordinate sets using ImageJ (NIH). The pre-scripted program then corrected every image tile by calling the ImageJ deformation function using that profile. Afterwards, we used the entire set of imaged tiles (full mouse brain in this case) to map out the tile-wise illumination profile. The images were grouped according to the stage movement, which affects the photo-bleaching profile. The program avoids using pixels that are considered empty background or dura artifacts using preset thresholds. Using those averaged profile tiles, the program normalized all the tile images. Please note, this profile is unique for each sample. Finally, the program picked 16 coronal slices (out of the nearly 2,000) with equal spacing and utilized ImageJ’s grid/collection stitching plugin to computationally stitch those 16 slides. The program then combined the transformation profiles from center to outer edge according to the calculated pairwise shifting distance. It used a tile-intensity weighted average to ensure the empty tiles did not contribute to the final profile. This approach significantly reduced the computational time and allowed parallelization with no communication overhead. The program automatically performed the aforementioned alignment and stitched the image set together. The program finally aligned the image sets if the sample was imaged through multiple runs during imaging acquisition.

### Computational: Vessel Digitization/Tracing

We started with interpolating the data into 1×1×1 μm resolution with cubic interpolation then subtracted the signal color channel (green) with the background color channel (red) to remove auto-fluorescent backgrounds. Next we performed a voxel binarization. The voxel with at least one of the following conditions passed as the foreground signal (vasculature), **a**. the voxel passed a fixed threshold (6x that of the non-empty space average) or **b**. passed a threshold (2.4x that of the non-empty space average) after subtracting a circular 35% local ranking filter. The binarized image was then skeletonized using 26-neighbor rule ^69^. The code then reconnected lose ends that were within 10 μm distance and removed all the short stem/furs shorter than 50 μm starting with the short ones and iterated until no more fur artifacts were found (Figure S1F). By using the binary image and the skeleton (center-line), the radius for each skeleton pixel can be measured. The code then grouped all the skeleton pixels into segments with the branching nodes, and all the segments shorter than 2x radius were further cleaned up with shortest graph path (Figure S1G). ROIs with poorly connected (<250 μm/node) were excluded in further analysis as shown Figure S1H. Finally, the code documented and traced all the segments and nodes with their connectivity, length, averaged radius, and raw skeleton locations. The full pipeline here is programed to be fully automatic and the code was fully vectorized and parallelized with reasonable memory consumption per thread (~8GB).

### Computational: Fluid Dynamic Simulation

The goal of calculating and visualizing a permeability tensor is to illustrate how well fluid can flow through the local microvasculature of a given volume in a given direction. Since the direction distribution of the microvasculature can be anisotropic, the fluid flow can move with a direction that is different from the pressure gradient direction, thus making the permeability in a tensor form. Such a tensor can illustrate the local microvascular performance and its directional characteristic.

The equation of permeability tensor is given by:

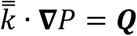

or

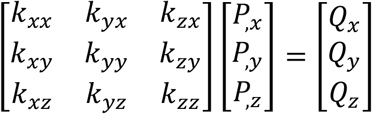

where *k* is the permeability tensor, *P* is the pressure, *Q* is the fluid flux, the subscript index is the Cartesian coordinate direction, and the comma is partial differentiation. We chose a size of 400 × 400 × 400 μm as the local representative control volume. We then probed the system with three ▽*P* that are the three unit-vectors in the Cartesian coordinate system. The pressure profile was applied on the surface of the cubical control volume, then the network flow profile was calculated by solving the system of equations of the Hagen–Poiseuille equation (with the viscosity set to unity for normalization) and conservation of flux. We chose the center cut plan to measure the directional flux and consequently, the permeability. Finally, to illustrate the vascular directionality of the isocortex, we projected the tensor onto penetrating, anterior-posterior, medial-lateral vectors according to their location within the isocortex using the equation 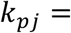 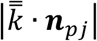, where subscript *pj* indicates the direction of the projecting vessels.

### Computational: Deep Learning Neural Network (DLNN) pericyte counting

We used a deep learning neural network (DLNN) to detect and classify cells. Instead of using a fully convoluted neural network like Unet, we chose to use per-cell multi-resolution-hybrid ResNet classification with potential cell locations. This makes the AI compute time significantly shorter. The potential cell locations were identified with local maximum within a radius of r = 8 μm. The image around the potential cell locations was fed to the network with two different resolutions. One is 101 x101 μm (101 × 101 pixel) and the other one is 501 × 501 μm (201 × 201 pixel). The two-window system allows the network to capture characteristics from two zoom scales simultaneously. In order to use global maximum at the end of the network, we stacked an empty (value zero) image onto the 3D direction of each image, which made them 101 × 101 × 2 and 201 × 201 × 2 pixels. We then assigned value 1 to the location of the potential cell, in this case, the center. At the end of the two networks of those two images, the intermediate images were flattened and concatenated into one. The classification was done with two bins, ‘pericytes’ and ‘everything else.’ The detailed schematic describing the network is in Figure S3.

We deployed two human annotators with the same training to annotate the data, and only used the mutually agreed data to train the AI to eliminate human error and bias. We used a strict set of criteria to include only capillary pericytes that have two subtypes (junctional and helical pericytes). Cells were counted only when the cell body was in the imaging plane and clear pericyte cell morphology could be detected. Cells associated with larger vessels, often with vascular smooth muscle morphology, were not counted to prevent the inclusion of erroneous cell types. We also excluded transitional cell types often referred to as either ensheathing pericytes or precapillary arteriolar smooth muscle cells, due to the controversy in the field as to whether this should be included as a pericyte subtype ^1,38^. A total 12,000 potential cell locations from multiple anatomical regions across 4 different brains were annotated by both annotators. 90% of the data selected at random was used to train the AI and the remaining 10% was used for validation. The 90% of the data taken for training was further truncated down to 3,400 potential cell locations with half positive and half negative for training. The positive cell selections in the raw data were around 19.6% (annotator #1) to 21.1% (annotator #2). The validation set was not truncated to represent actual performance. The performance can be found in Table S3.

### Computational: Deep Learning Neural Network (DLNN) nNOS neuron counting

The morphology and size of tdTomato positive cells in the granular layer of the cerebellum from nNOS-CreER:Ai14 mice differs significantly from other tdTomato positive nNOS neurons in other brain regions. Thus, we developed new DLNN AI algorithms to consider not only cell morphology but also the location of cells by putting additional zoomed-out, low-resolution images of whole coronal sections. The network set-up is similar to the pericyte classification with one more image containing the coronal section with the cell location. The inputs are 101 ×101 μm (101 × 101 pixel), 501 × 501 μm (201 × 201 pixel), and the full frame low resolution 12 × 8 mm (201 × 201 pixels). Similar to the pericyte network, we made images 101 × 101 × 2, 202 × 202 × 2, and 202 × 202 × 2 pixels with the cell location marked as value 1. At the end, those three sub-networks were flattened and concatenated into one. The classification is done with three bins, nNOS neurons, cerebellar granular nNOS neurons, and everything else. One human user created 10,000 annotations from 5 pan nNOS and 5 nNOS subtype brains. 5,000 cells from 5 brains were initially used to train the AI. Another 5,000 cells from 5 new brains were used to evaluate the AI performance. The AI reached an F1 score = 0.96, which is comparable to human performance. The details for the network are in Figure S3. The performance can be found in Table S3.

### Isocortical flatmap

We started with Allen CCF annotation images to solve the Laplace equation by setting the surface of cortical layer 1 as potential ‘1,’ the surface of layer 6b as ‘0,’ and the surface of everything else as flux ‘0’ ^31^. We used the potential map to find the gradient direction as the projecting direction. The projection was first traced to the cortical surface and then flattened at the Anterior-Posterior (A-P) tangential plane, which later preserved the A-P coordinate on the flat map. The flattened map has the y-axis mapped as the original A-P coordinate at the surface, and the x-axis was adjusted to represent the surface arc (azimuth) length to the reference *X*-zero. The reference *X*-zero was defined on the cortical ridge in the dorsal direction (maximum *Y* point in 3D) with a straight cut in the A-P direction. Finally, the projection profile was saved at two resolutions, 10 × 10 × 10 μm^3^ and 20 × 20 × 20 μm^3^. We created a Matlab script that can map any signal (previously registered to the Allen CCF) into a 3D projected isocortical flatmap.

### Conversion of 2D based counting to 3D cell density

STPT imaging has very accurate cutting and stage depth movement, which allows us to convert the 2D cell counting to 3D cell density. We used previously calculated 3D conversion factors for cytoplasmic (factor = 1.4) and nuclear signals (factor = 1.5) to generate density estimates of nNOS neurons and other neuronal cell type datasets ^20^. To estimate the 3D conversion factor for pericytes as we have done for Figure S1 from ^20^, we imaged one PDGFRβ-Cre:Ai14 mouse brain with 1 × 1 × 5 μm, as done with vascular imaging (Figure 1B). Then, we cropped out 40 ROIs with 500 × 500 × 50 (x,y,z) μm^3^ in size randomly from different areas including the cortex, hippocampus, midbrain, hypothalamus, and cerebellum. We then manually counted pericytes in 2D (5^th^ z slice from the stack) and 3D (total 10 z slices from the stack). We counted total 840 cells from 2D counting and 1769 cells from 3D counting (3D/2D ratio = 2.13 ± 0.28, mean ± standard deviation), resulting in 3D conversion factor of 2.1, which was applied as a conversion factor to estimate pericyte numbers in 3D.

To estimate the anatomical volume from each sample, the Allen CCF was registered to individual samples first using Elastix ^70^. Anatomical labels were transformed based on the registration parameters and the number of voxels associated with specific anatomical IDs were used to estimate the 3D volume of each anatomical area ^20^.

### In vivo two-photon recording and comparison with STPT vascular measurement

#### Surgery

All surgeries were performed under isoflurane anesthesia (in oxygen, 5% for induction and 1.5-2% for maintenance). A custom-machined titanium head bolt was attached to the skull with cyanoacrylate glue (#32002, Vibra-tite). The head bolt was positioned along the midline and just posterior to the lambda cranial suture. Two self-tapping 3/32” #000 screws (J.I. Morris) were implanted into the skull contralateral to the measurement sites over the frontal lobe and parietal lobe. For measurements using two-photon laser scanning microscopy (2PLSM), a polished and reinforced thin-skull (PoRTS) window was made covering the right somatosensory cortex as described previously ^15,71^. Following the surgery, mice were returned to their home cage for recovery for at least one week, and then started habituation on experimental apparatus. Habituation sessions were performed 2-4 times over the course of one week, with the duration increasing from 5 min to 45 min.

#### Measurements using two-photon laser scanning microscopy (2PLSM)

Mice were briefly anesthetized with isoflurane (5% in oxygen) and retro-orbitally injected with 50 μL 5% (weight/volume in saline) fluorescein-conjugated dextran (70 kDa, Sigma-Aldrich, cat.no.: 46945), and then fixed on a spherical treadmill. Imaging was done on a Sutter Movable Objective Microscope with a 20X, 1.0 NA water dipping objective (Olympus, XLUMPlanFLN). A MaiTai HP (Spectra-Physics, Santa Clara, CA) laser tuned to 800 nm was used for fluorophore excitation. All imaging with the water-immersion lens was done with room temperature distilled water between the PoRTS window and the objective. All the 2PLSM measurements were started at least 20 minutes after isoflurane exposure to reduce the disruption of physiological signals due to anesthetics. High-resolution image stacks of the vasculature were collected across a 500 by 500 μm field and up to a depth of 250 um from the pial surface. All the images were acquired with increasing laser power up to 100 mW at a depth of ~200 μm. Lateral sampling was 0.64 um per pixel and axial sampling was at 1 um steps between frames. Shortly (within 20 minutes) after the imaging, the mouse was perfused with FITC filling for STPT based *ex vivo* vasculature imaging.

### In vivo and ex vivo comparison

In order to compare our measurements for vessel radii in STPT imaging datasets to vessel parameters measured *in vivo*, the same animals that were used for 2PLSM (See **In vivo two-photon recording and comparison with STPT vascular measurement**) underwent the FITC-fill perfusion and STPT imaging steps described above. However, STPT imaging was only conducted on the cortical hemisphere used for 2PLSM, with imaging spanning from prefrontal regions to visual cortex regions, in order to appropriately capture the primary somatosensory cortex limb region. Following stitching and tracing of the images, the raw imaging data was reconstructed in 3D in order to visualize the cortical surface. To find the 2PLSM imaging window, vessel landmarks used for navigation purpose in 2PLSM were again used to identify the same landmark vessels in the STPT imaging dataset (Figure S2). The region of interest was further confirmed by anatomical landmarks (proximity to bregma, surface vessels, etc.) through overlay of STPT and 2PLSM imaging window regions. Next, within a STPT imaging z stack, borders were inserted using ImageJ software to further outline the in vivo imaging window region in the 3D data. Then the in vivo imaging z stack data were used to identify branch points along the penetrating vessel tracked during 2PLSM. This provided identifiable characteristics to further locate the same vessel in the STPT imaging dataset. Once the exact vessel was identified in the STPT images, the precise 3D coordinates were tracked to accurately obtain the radii measurements from the traced vessel data, see the **Computational: Vessel Digitization/Tracing section** for details. In 2PLSM images, vessel diameter measurements were manually taken with adjusted pixel/micron distances using the straight-line function in ImageJ. These vessel diameter measurements accounted for the lumen of the vessel, at half of the maximum fluorescence intensity profile and were adjusted for pixilation of 2PLSM data. These measurements have been further refined through VasoMetrics ImageJ macro ^72^. To identify the radii and diameter measurements from the STPT imaging data, exact vessel coordinates were used to retrieve the associated vessel radii measurements using custom MATLAB code. A minimum of 10 vessel diameter measurements were taken per imaging window (each animal contained 2 imaging regions of interest) per animal.

### Statistical analysis

All statistical analysis, including multi-region of interest (ROI) correlation analysis, was done in Matlab (Mathworks). We used an averaged value of the experimented animals while treating each ROI as an individual data point to calculate the correlation coefficient *R* between vascular and cell density measurements. The p value was calculated based on the null hypothesis that the two groups have no correlation; the values were adjusted with the Bonferroni correction for multiple comparison correction.

## Acknowledgments

This publication was made possible by an NIH grant R01NS108407 to Y.K., R01NS078168 and R01NS101353 to P.J.D. Its contents are solely the responsibility of the authors and do not necessarily represent the views of the funding agency. We thank Drs. Nanyin Zhang, Anirban Paul, Byungkook Lim for construction discussion of the manuscript, Dr. Volkhard Linder for kindly sharing PDGFRβ-Cre transgenic mice and Rebecca Betty for assistance in editing the manuscript. We acknowledge use of computational resources in the High Performance Computing cluster at the Penn State College of Medicine.

## Contributions

Conceptualization, Y.K.; Data Collection, H.C.B, U.C., Y.K.; Developing Computational Analysis, Y.W.; Data Analysis, Y.W., H.C.B., U.C.; In vivo two-photon imaging, Q.Z., P.J.D.; Neuronal subtypes STPT data collection; R.M., P.O.; Web visualization, D.J.V., K.C.; Manuscript preparation: Y.K., Y.W., H.C.B. with help from other authors.

## Data Availability

High-resolution serial two-photon tomography images can be found at https://kimlab.io/brain-map/nvu/

We deposit all full resolution image datasets in the Brain Image Library (https://www.brainimagelibrary.org/). We plan to supply the download link before full publication.

Simulation-ready dataset for cerebrovascular tracing can be downloadable at https://kimlab.io/data_share/files/NVU_young/vessel_young_simulation-ready.7z

Custom-built codes including isocortical flatmaps are available as Supplementary data (Code S1-S3)

All dataset and codes can be used for non-profit research without any restriction.

