## Supplementary Info for "The cellular architecture of microvessels, pericytes and neuronal cell types in organizing regional brain energy homeostasis in mice"

### Supplementary information

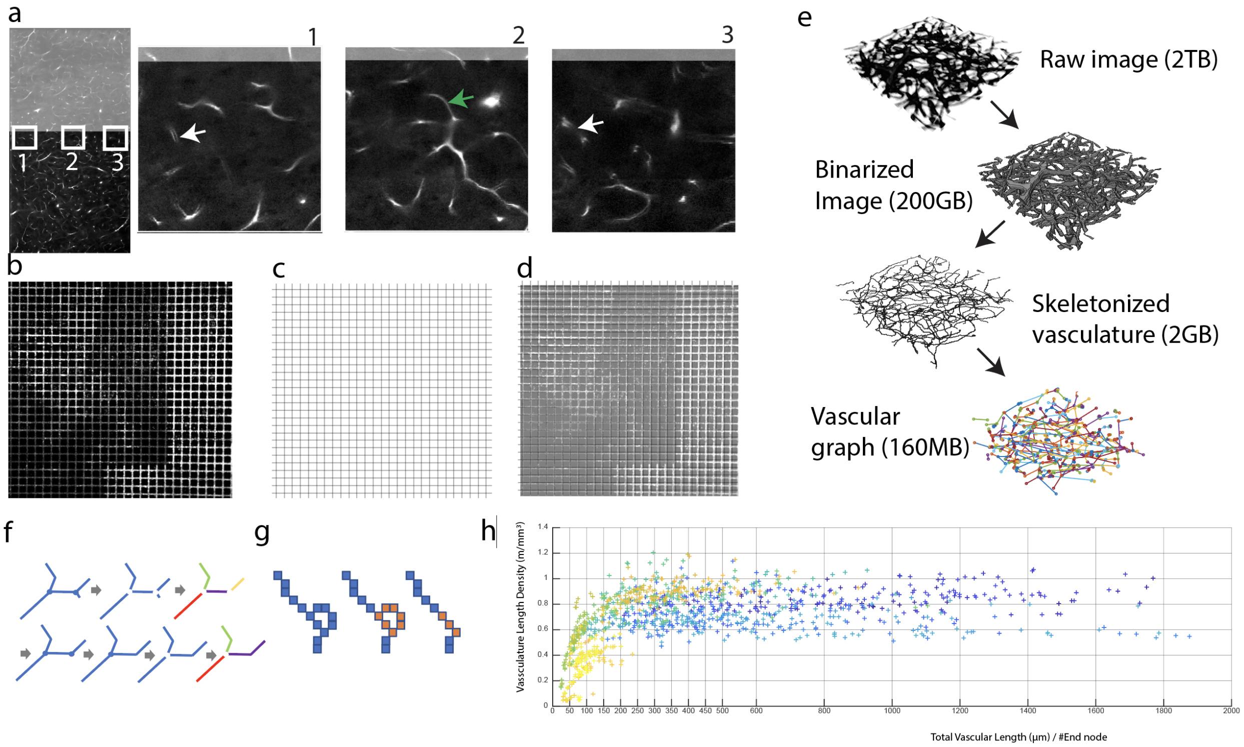


**Figure S1. Stitching correction, data processing pipeline, and quality control for vascular imaging**

**a**. Correcting for optical aberration from an objective lens that impacts the vasculature tracing in two adjacent image tiles. When we manually aligned two adjacent tiles to match the middle part of the tile (e.g., a green arrow in area 2), vascular signals in the left (area 1) and right (area 3) side of tiles did not align (white arrows). **b**. The EM-grid imaged under the STPT system. The EM grid is a known orthogonal grid line, serving as a ground truth. Thus, any distortion from STPT imaged EM grid can help to assess optical aberrations introduced by the scope. **c**. A theoretical perfect gridline. **d**. Overlap of (**b**) and (**c**). Matching grid points between (**b**) and (**c**) were marked and used to calculate the deformation profile for aberration correction in our stitching algorithm. **e**. Data size throughout the analysis pipeline. **f-g**. Schematic shows how the algorithm groups, truncates the short branching, and straightens the looping artifact. The straightening showing in (**g**) is through shortest path. Those artifacts nodes that cannot be resolved this way were replaced with center representative pixel-equivalents for the later computation. The final network was processed with six such iterations. **h**. Each data point represents a ROI in one mouse. The plot includes all the animal data to check tracing quality control. The x axis is total vascular length divided by the number of end node. ROIs in the lower value in the x axis have higher probability to include higher number of dysconnectivity due to imaging artifact. The color of each data point is according with Allen ontology. We picked 250 as the threshold since the dysconnectivity no longer affected the length density after passing 250. All data points that did not pass this threshold were excluded in our final analysis.

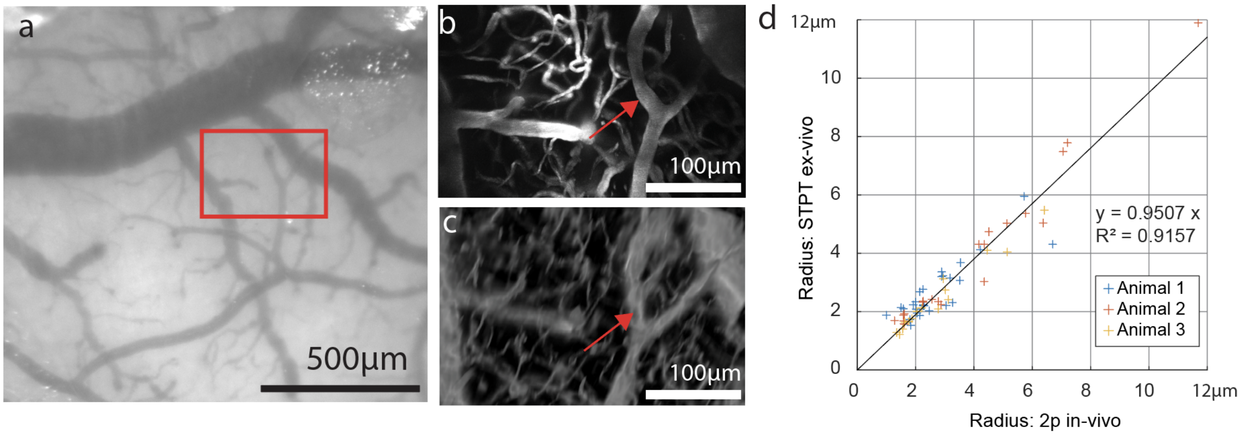


**Figure S2. *In vivo* and *ex vivo* comparison.**
**a**. A zoom out view of the brain pial vessels for two-photon imaging of the areas of interest (**b**). **b**. Maximum intensity projection (200 µm thick) of *in vivo* two-photon imaging from the red boxed area. **c**. The corresponding region in the ex-vivo STPT image volume (translated to match the imaging angle of the *in vivo* imaging). Red arrows in (**b** and **c**) indicates the same Y shaped branch identified in both imaging modalities to demonstrate preserved vascular architecture in the STPT imaging. **d**. Radius measurements from same vessels identified in both *in vivo* and *ex vivo* imaging (n=3 animals) showed near identical results.

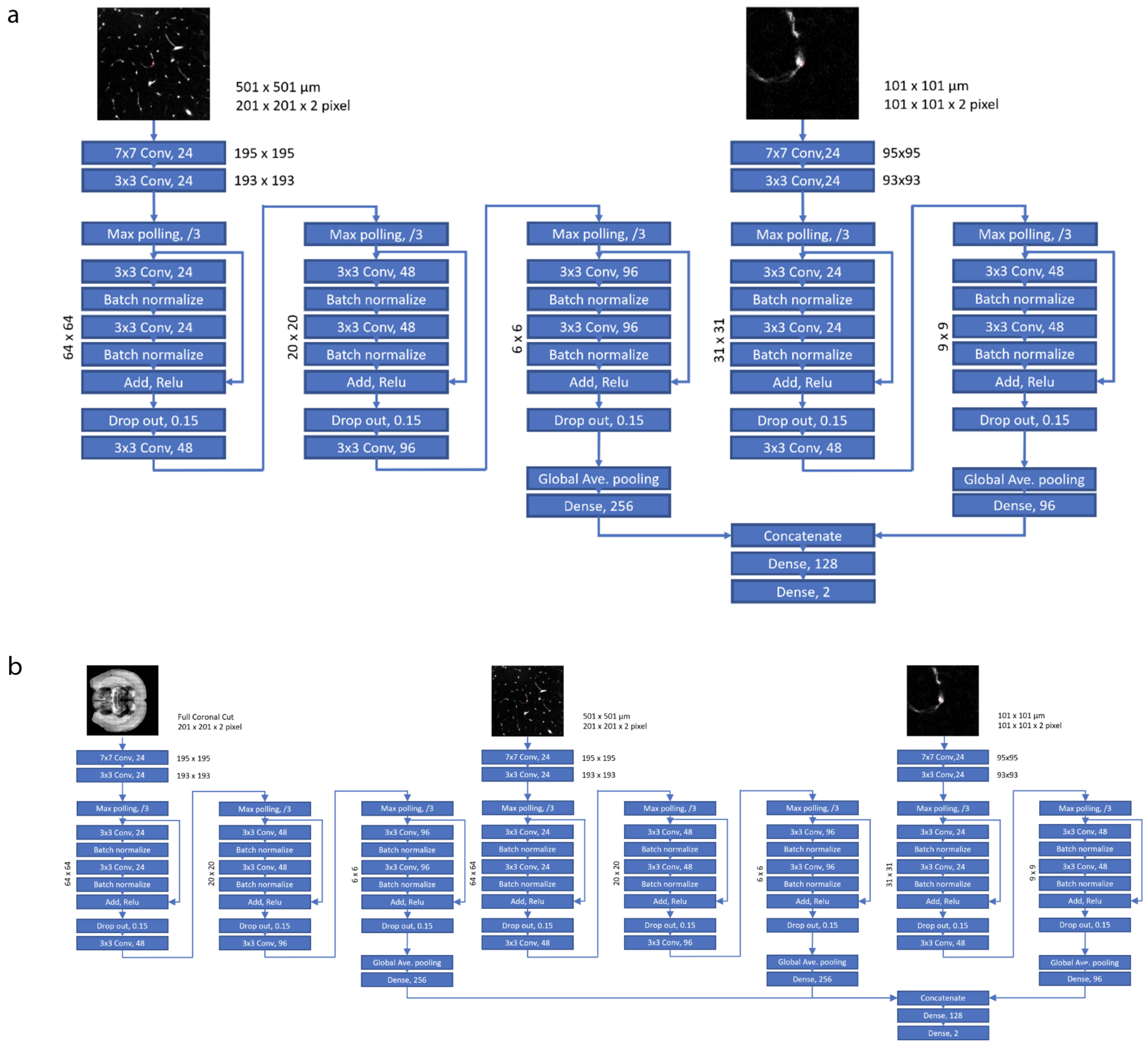


**Figure S3. Deep Learning Neural Network for pericyte and nNOS cell counting**
**a**. The network used in pericyte counting. **b**. The network used in nNOS and its subtype counting. Both networks are adapted version of RESNET with concatenating of multiple resolution images to bring the awareness of the nature of the cell to the network; Such as its shape in high resolution, the neighboring background feature in mid resolution, and the brain location in the low resolution.

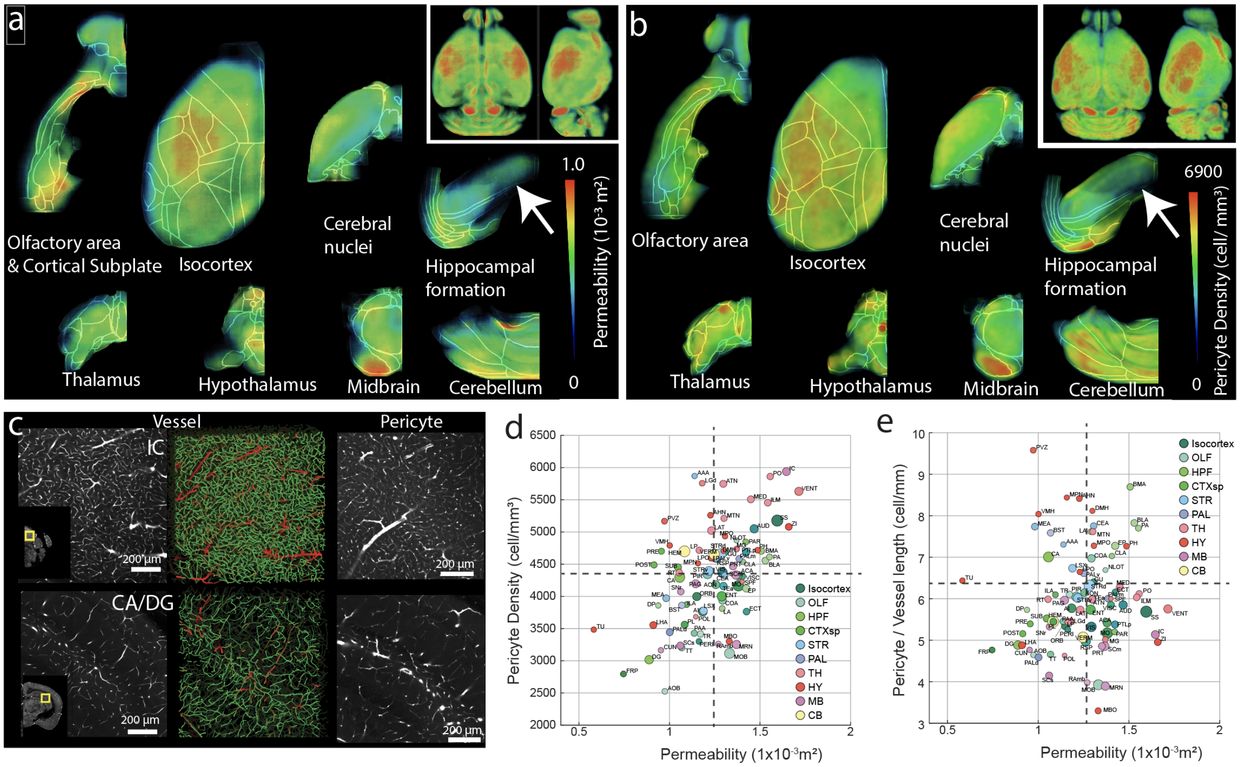


**Figure S4. Low vascular permeability and pericyte densities in the hippocampal areas.** **a-b**. Graphical representation of spherically integrated permeability results (**a**) and pericyte density (**b**) of the whole brain. White arrows highlight relatively low vascular permeability and pericyte densities in the hippocampal areas. See also Table S3 and S6. **c**. Examples of high densities of vasculature and pericytes in the inferior colliculus (IC) and Ammon's horn/dentate gyrus (CA/DG) of the hippocampus. Mid panels represent detected vasculature in 100 µm thick sections (green: microvessels, red: large vessels). **d-e**. Scatter plot of pericyte and vascular permeability (**d**), and pericyte vascular coverage (pericyte/vessel length) and vascular permeability.

**Table S1. Vascular analysis data**

This table documents all vascular length (vess_length, m), vascular length density (vess_length, m/mm$^3$), averaged vasculature radius (vess_radii, µm), vasculature branching density (vess_branching, 1/mm$^3$), and total volume of each region of interest (ROI_volume, mm$^3$). 'Nan' in the table indicating the quality of the ROI did not pass the connectivity quality control and was removed.

**Table S2. The performance metric of the multi-resolution DLNN counting.**

Both the pericyte counting AI and the neuron counting AI was benchmarked against two expert annotators. In pericyte AI, we also show the performance improvement of using human agreed data in training.

**Table S3. Pericyte density**

Density unit of the pericyte is 1/mm$^3$.

**Table S4. nNOS and nNOS subtype density**

3D density of nNOS (nnos), nNOS co-expressing Neuropeptide Y (nnos_npy), Somatostatin (nnos_sst), Parvalbumin (nnos_pv), or Vasoactive Intestinal Peptide (nnos_vip). Density measurement unit is 1/mm$^3$.

**Table S5. Glutamatergic and GABAergic cell type density**

3D density of parvalbumin (PV), somatostatin (SST), vasoactive intestinal peptide (VIP), pan GABAergic (Gad2), and pan-glutamatergic (vGlut1) neurons. Density measurement unit is 1/mm$^3$.

**Table S6. Computationally simulated permeability results**

This table documented the simulated permeability (m$^2$).'Nan' in the table indicating the quality of the ROI did not pass the connectivity quality control and was removed.

**Table S7. Vascular length density and pericyte density in the dorsal striatum**

Vascular length density (vess_length, m/mm$^3$) and pericyte density (cell/mm$^3$) in subregions of the dorsal striatum.

**Movie S1. Cerebral vascular measurements**

Top left: Allen CCF with anatomical segmentation lines overlaid. Top right: Vessel length density. Bottom left: Radius measurement. Bottom right: Vascular branching density. Numbers in the top left corner represent estimated A/P bregma coordinates.

**Movie S2. Pericyte density**

Left: Allen CCF with anatomical segmentation lines overlaid. Right: Pericyte density. Numbers in the top left corner represent estimated A/P bregma coordinates.

**Movie S3. nNOS subtypes density**

Top left: Allen CCF with anatomical segmentation lines overlaid. Top middle: nNOS neuron density. Top right: density of nNOS co-expressing NPY. Bottom left: density of nNOS co-expressing PV. Bottom middle: density of nNOS co-expressing SST. Bottom right: density of nNOS co-expressing VIP. Numbers in the top left corner represent estimated A/P bregma coordinates.

**Movie S4. Neuronal subtypes density**
Top left: Allen CCF with anatomical segmentation lines overlaid. Top middle: vGlut1 neuron density. Top right: Gad2 neuron density. Bottom left: PV neuron density. Bottom middle: SST neuron density. Bottom right: VIP neuron density. Numbers in the top left corner represent estimated A/P bregma coordinates.

**Movie S5. Vascular permeability simulation results.**
Left: Allen CCF with anatomical segmentation lines overlaid. Right: Simulated permeability results. Numbers in the top left corner represent estimated A/P bregma coordinates.

**Code S1. STPT Vasculature Tracing and Analysis**
This package contains the code used in vasculature tracing and analyzing. It includes a stitching algorithm to correct all the intensity and abberation to make the image tracible, a tracing pipeline to translate the STPT images into documented vasculature centerlines/nodes, an analysis panel to digest the results into Allen CCF ontology, and finally a simulation solver to find all the permeability tensor voxels from the traced data. Please refer to the README.md in the package for more information.

**Code S2. Cortical Flat Map**
This package contains the code for producing cortical flat map form any images that are registered/transformed onto the Allen CCF. Please refer to the README.md in the package for more information.

**Code S3. Multi Resolution DLNN Cell Counting**
This package contains the cell counting pipeline used in this paper. It includes the GUI tools for creating ground truth annotations, the network in Tensorflow 2.0, the training protocols, and a pipeline to digest the full brain data into counted cells in Allen CCF ontology. Please refer to the README.md in the package for more information.
